# Discovery of 20 novel ribosomal leaders in bacteria and archaea

**DOI:** 10.1101/2020.01.29.924993

**Authors:** Iris Eckert, Zasha Weinberg

## Abstract

**Background:** RNAs perform many functions in addition to supplying coding templates, such as binding proteins. RNA-protein interactions are important in multiple processes in all domains of life, and the discovery of additional protein-binding RNAs expands the scope for studying such interactions. To find such RNAs, we exploited a form of ribosomal regulation. Ribosome biosynthesis must be tightly regulated to ensure that concentrations of rRNAs and ribosomal proteins (r-proteins) match. One regulatory mechanism is a ribosomal leader (r-leader), which is a domain in the 5′ UTR of an mRNA whose genes encode r-proteins. When the concentration of one of these r-proteins is high, the protein binds the r-leader in its own mRNA, reducing gene expression and thus protein concentrations. To date, 35 types of r-leaders have been validated or predicted.

**Results:** By analyzing additional conserved RNA structures on a multi-genome scale, we identified 20 novel r-leader structures. Surprisingly, these included new r-leaders in the highly studied organisms *Escherichia coli* and *Bacillus subtilis*. Our results reveal several cases where multiple unrelated RNA structures likely bind the same r-protein ligand, and uncover previously unknown r-protein ligands. Each r-leader consistently occurs upstream of r-protein genes, suggesting a regulatory function. That the predicted r-leaders function as RNAs is supported by evolutionary correlations in the nucleotide sequences that are characteristic of a conserved RNA secondary structure. The r-leader predictions are also consistent with the locations of experimentally determined transcription start sites.

**Conclusions:** This work increases the number of known or predicted r-leader structures by more than 50%, providing additional opportunities to study structural and evolutionary aspects of RNA-protein interactions. These results provide a starting point for detailed experimental studies.

## Background

The ribosome is an RNA-protein complex that performs protein synthesis in all living cells [1–3]. The ribosome consists of two subunits: the small subunit binds the mRNA template, while the large subunit catalyzes the peptidyl transfer reaction. Each bacterial or archaeal ribosome is made of three different rRNAs (5S, 16S and 23S) and many ribosomal proteins (r-proteins). Ribosome synthesis is a complex process involving multiple maturation steps, including the processing and folding of rRNA through the binding of r-proteins.

Because of the central importance of the ribosome to cellular function, ribosomes consume a large portion of the cell’s energy [4]. As a result of this huge cost in energy, cells use highly optimized regulatory systems to ensure that r-proteins are at their optimal concentrations [4].

One common regulatory system in bacteria is a type of feedback regulation known as a ribosomal leader (r-leader) [4–8]. R-leaders are structured RNA elements that occur in the 5′ UTRs of mRNAs whose genes encode r-proteins. One or more of these r-proteins can interact with the r-leader, in addition to the r-protein’s normal role in the ribosome. Excess r-proteins bind the r-leader, which leads to a change in the 5′ UTR’s secondary structure that results in repressed expression of the downstream genes. Known mechanisms for repression [7] include the conditional formation of Rho-independent transcription terminators and sequestration of the ribosome-binding site.

Because the r-protein ligands of r-leaders also bind rRNA, it was hypothesized that r-leaders imitate the structure of the relevant rRNA binding site [5, 7]. In several cases, this similarity is apparent from the secondary structure [9–11], while other cases require a crystal structure to demonstrate structural mimicry [12, 13]. However, it is not a requirement that the r-leader must copy the rRNA structure; in a few cases, no meaningful similarities could be detected [7].

Thirty-five r-leaders have been confirmed or proposed (Additional file 1: Table S1) [7]. Knowledge of r-leaders is important to provide a complete picture of ribosome assembly and overall cellular metabolism.

R-leaders also provide a starting point for deeper research into the structure and evolution of RNA-protein interactions [14], which are important in all domains of life in a variety of contexts. For example, some r-proteins are the ligand of multiple types of r-leaders that each have different conserved sequence and structural features. These distinct structures thus exhibit multiple solutions to a single biochemical problem, and create opportunities [15] to study the evolutionary and structural influences leading to these end-points.

Additionally, r-leaders can be a model system to understand *cis*-regulatory mechanisms. For example, only a handful of *cis*-regulatory RNAs in archaea are experimentally confirmed or even predicted [16, 17]. Information on this aspect of gene regulation is therefore lacking.

We therefore decided to detect novel r-leaders using a bioinformatics strategy centered around a phenomenon known as covariation. Covariation typically refers to mutations in which both nucleotides involved in a Watson-Crick base pair change and the resulting base pair is also energetically favorable. With sufficient evolutionary time, such mutations are frequent in RNAs that conserve a structure, but only occur sporadically by chance in sequences that do not function as RNAs. Analysis of covariation has been highly successful in determining conserved structures of molecules, such as ribosomal RNAs, that were later confirmed experimentally [18–20]. Covariation-based strategies have also been used to find RNAs *de novo*, which have resulted in experimentally validated riboswitches [21], ribozymes [22] and an r-leader [23, 24], among others.

## Results

### Discovery and evaluation of candidate r-leaders

To find novel r-leaders, we inspected raw computer predictions of conserved RNA secondary structures from earlier studies [25, 26]. Each prediction consisted of a multiple sequence alignment and a conserved secondary structure. We call these predictions “motifs”. In searching for r-leaders, the most promising motifs are those that are frequently located upstream of genes that encode r-proteins, and are thus potential *cis*-regulators of these genes. We analyzed the 32 best predictions by finding additional homologs and exploiting covariation to find additional conserved RNA secondary structure, using previously established approaches [25–27].

After this analysis, motifs whose secondary structures were supported by covariation and that remained consistently upstream of genes encoding r-proteins were considered candidate r-leaders. The evaluation of covariation is often not straightforward [26] (see Methods). To ensure that none of our motifs were previously discovered, we compared them to previously established RNAs. We compiled a list of all r-leaders that are experimentally verified or have a predicted alignment (Additional file 1: Table S1). We eliminated candidate motifs whose homologs overlap previously published RNAs, and those whose primary and secondary structure features were essentially the same as any previously published r-leader (see Methods). However, we included a new S4 r-leader in Fusobacteria whose potential binding site resembles that of the previously published Firmicutes S4 r-leader, but occurs in the context of a different secondary structure. We thus report 20 novel r-leader motifs (Table 1). The 20 new r-leaders show no meaningful similarities to one another (see Methods), except for a partial resemblance between the archaeal S15 leaders (see below). We refer to the new motifs (Table 1) based on their most likely ligands and the lineage of bacteria or archaea in which they occur (e.g., the Fusobacteria S4 r-leader motif).

**Table 1.**
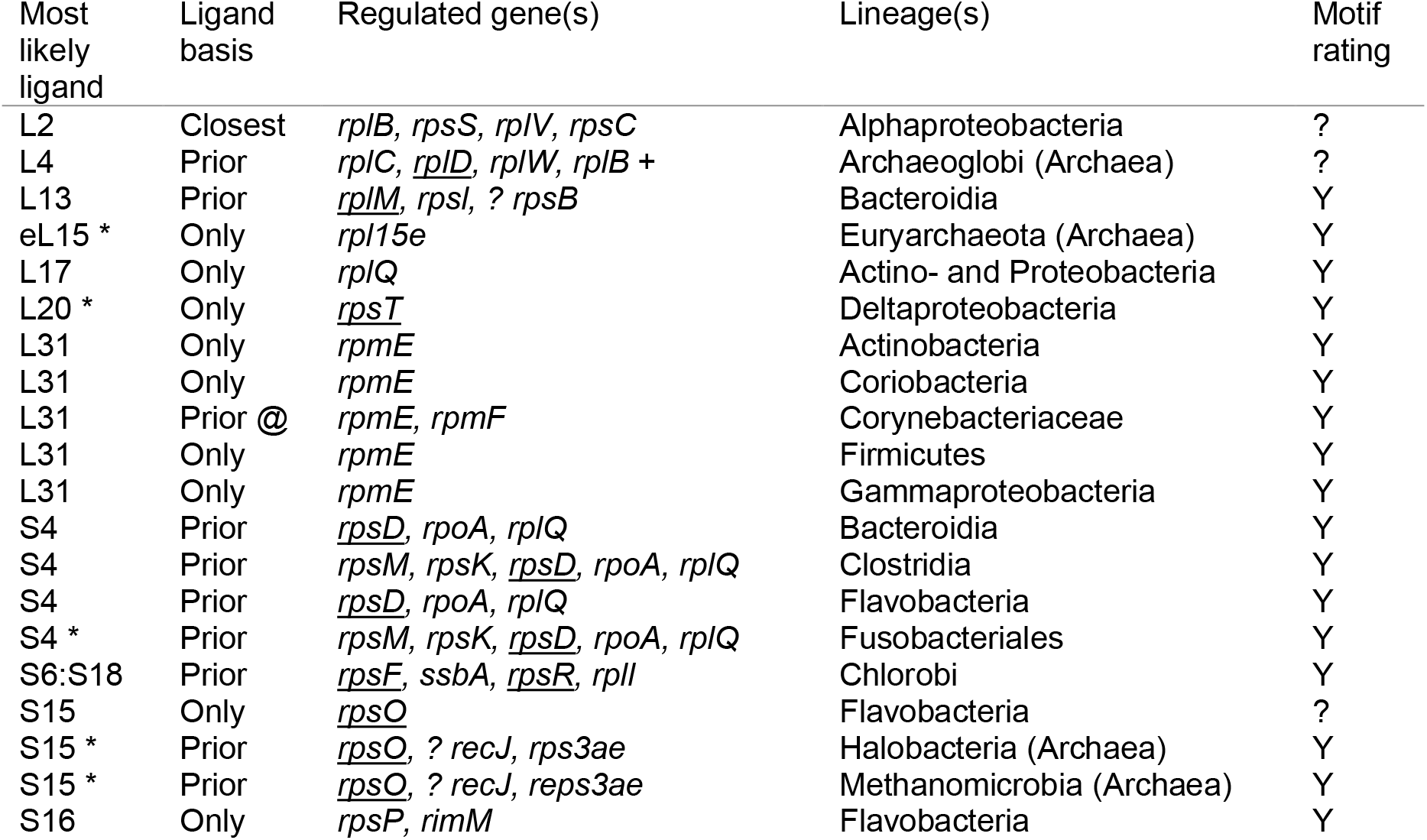
Summary of novel ribosomal leaders. “Most-likely ligand”: see text. All proteins are the bacterial version, except eL15 is the eukaryotic/archaeal “L15” protein. Some principles used for ligand predictions are uncertain (see text). Asterisk (*): r-leader is potentially similar to rRNA binding site for the given ligand, supporting its assignment. “Ligand basis”: principle with which the most likely gene was hypothesized. “Only”: there is only one regulated r-gene; “Prior”: there are multiple regulated genes and one encodes a known ligand of another r-leader (at sign (@): based on L31 motifs in the current paper), “Closest”: the immediately downstream gene. “Regulated gene(s)”: genes consistently located in potentially regulated operons are listed in the order they most often appear. Most genes encode ribosomal proteins. Genes encoding the ligand of a previously established r-leader are underlined. Motifs lacking underlined genes bind novel r-protein ligands. Question mark: possible operon end. Plus sign: apparent operon can often be extended. “Lineage”: taxon containing the motif. Archaeal taxa are indicated. “Rating”: covariation evidence supporting assignment as RNA. “Y” = likely candidate, “?” = borderline candidate. The reasons for assigning candidates as borderline are given in Additional file 1: Supplementary text.

In addition to the 20 novel r-leaders, we found three motifs (Additional file 1: Figure S1) whose secondary structure and nucleotide conservation patterns are fundamentally the same as already-established motifs, but that included additional homologs not previously found. These were the L25 r-leader in Enterobacteria [28], the S10 r-leader in Firmicutes [29], where we found numerous examples in the sub-lineage Clostridia and the L19 leader in Firmicutes [29], where we found many examples in the distinct phylum Flavobacteria. These alignments are available as Additional files (see below), but are not further discussed in this manuscript.

All novel motifs are summarized in Table 1. Statistics regarding base pairs and covariation are provided (Additional file 1: Table S2), and include statistically significant covariation signals (see Methods) for all 20 r-leader motifs. For each motif, we also provide alignments and information on downstream genes and taxonomy (Additional file 2) and machine-readable alignments with (Additional file 3) and without (Additional file 4) additional metadata. We found some stems with very borderline support from covariation. These stems are indicated only in Additional files 2 and 3. Consensus diagrams for all novel motifs, previously published motifs that bind the same putative ligand, as well as the relevant protein-binding sites in rRNAs are found in Additional file 1: Figures S2-S11.

### Transcriptome data

Each of the 20 r-leader motifs has between 67 and 8,743 examples in various genomic locations in various organisms, with a total of 29,730 r-leader examples (Additional file 1: Table S2). We hypothesize that each r-leader example regulates its immediately downstream gene, and possibly additional co-transcribed genes. Thus, if our hypothesis is correct, each r-leader example should be located downstream of the transcription start site (TSS) for its regulated gene. By contrast, if the TSS occurs downstream of the r-leader, the r-leader would not be transcribed, which could suggest that our hypothesis is incorrect. A TSS could also be located within the r-leader such that important conserved features would not be transcribed. Such a TSS position would also contradict our hypothesis. Therefore, we wished to determine if TSS positions are consistent with our r-leader predictions.

TSS positions could, in principle, be determined on a genome-wide scale using standard RNA-seq experiments. However, such experiments do not provide reliable TSS predictions, because an apparent TSS in RNA-seq data could correspond to the 5′ end of a processed RNA [30]. Fortunately, differential RNA-seq (dRNA-seq) addresses this problem [30]. We thus searched (Additional file 1: Supplementary text) for studies that used dRNA-seq or related methods to determine TSS sites for organisms that contain a predicted r-leader. In some cases, no TSS was provided for the gene that we predicted is regulated by the relevant r-leader. We did not analyze these cases, since we cannot be certain where the relevant TSS is.

Ultimately, we found TSS positions for 11 distinct regulated genes out of the 29,730 r-leader examples (Fig. 1, Additional file 1: Table S3). These 11 regulated genes include examples of eight of the 20 motifs. (Some motifs have multiple examples with TSS data.) Most organisms lack published dRNA-seq results, and most of our motif examples occur in metagenomic sequences, which also lack dRNA-seq experiments. Therefore, only a small fraction of the total r-leader examples had usable TSS data, and we analyzed these available data.

**Figure 1.**
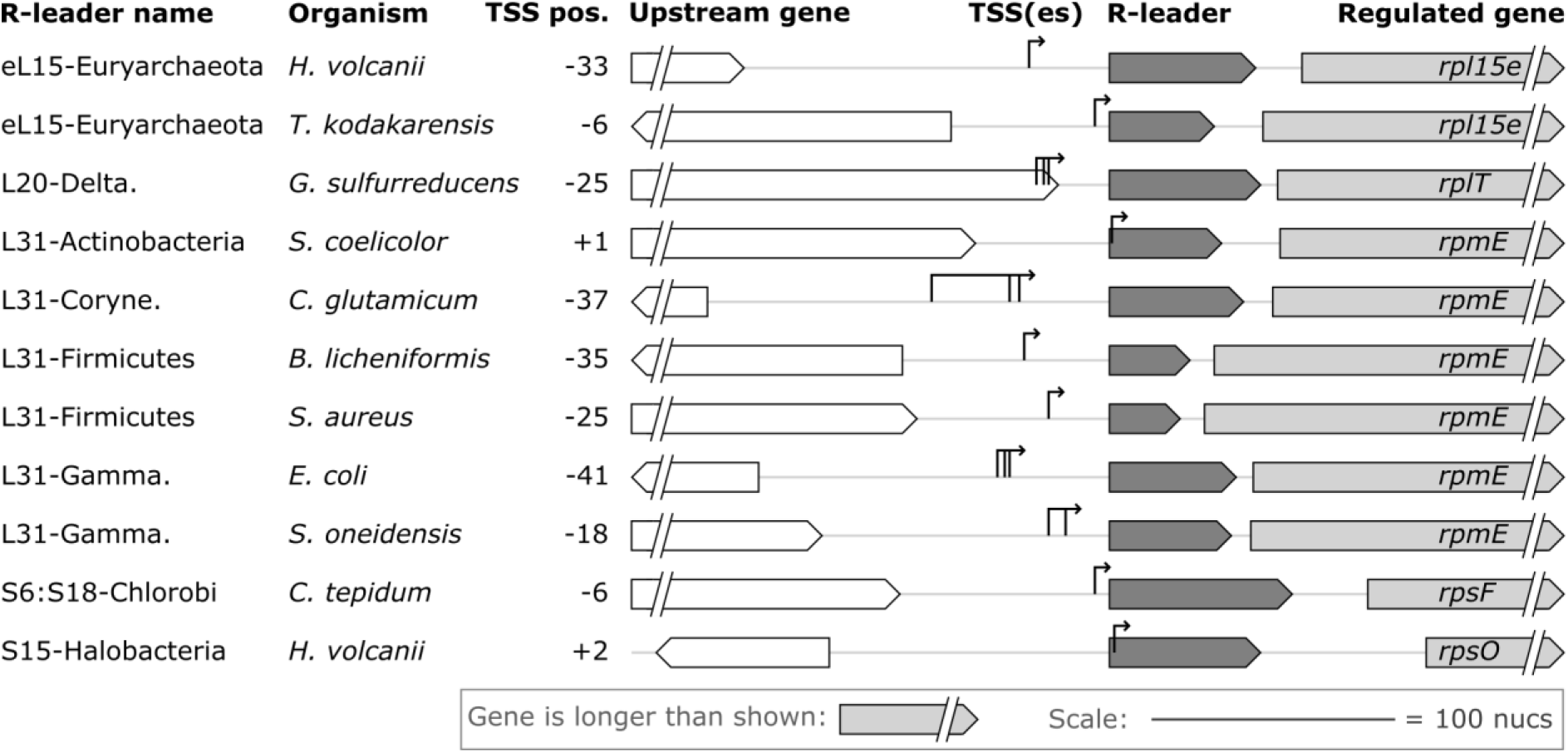
Experimentally determined transcription start sites (TSSes) are consistent with a *cis*-regulatory role for our r-leader motifs. We found experimentally annotated TSS data for 11 specific genes that we predicted to be regulated by an example of one of the 20 new r-leaders (see text). “R-leader name”: predicted ligand and lineage of relevant r-leader (refers to Table 1). Some lineages are abbreviated. “Organism”: the organism in which TSS experiments were conducted. “TSS pos.”: position of the TSS relative to the 5′ end of the r-leader. If there are multiple TSSes, the TSS nearest the gene is used. Negative values mean that the TSS is upstream of the r-leader, so the r-leader is transcribed. Positive values mean that the given number of r-leader nucleotides would be skipped. These numbers are low (at most +2; see text). Genome cartoons are to scale (scale bar: lower right). Genes and r-leaders are depicted as arrows whose direction corresponds to their DNA strand. The TSS or TSSes for the putatively regulated gene are shown as vertical lines, with a thin arrow above them. The name of the regulated gene is given. Additional data, citations and underlying numbers are provided (Additional file 1: Table S3).

As expected, the TSS sites for all 11 genes were positioned so as to transcribe the r-leader (Fig. 1). In two cases, there were TSSes that would cause up to two nucleotides of the r-leader to be skipped (Fig. 1, see “L31-Actinobacteria” and “S15-Halobacteria”). However, in both cases, the skipped nucleotides are not well conserved, and therefore likely do not belong to the r-leader. Thus, in all cases, conserved nucleotides and stems are located downstream of all TSSes

### Analysis of ligands and binding sites

To generate hypotheses for the r-protein ligands of the motifs, we assumed that the ligand would be encoded by a gene regulated by the r-leader, as this is true in all known cases. Eight novel r-leader motifs were observed only upstream of genes encoding one r-protein (Table 1), and in these cases we concluded that this r-protein is the ligand. A ninth motif is directly upstream of *rpsP*, which encodes r-protein S16, and also usually upstream of *rimM* genes, which are involved in 16S rRNA processing. We believe this motif likely functions as an S16 r-leader (Table 1) despite the *rimM* genes (explained in Additional file 1: Supplementary text, subheading “S16”).

The remaining 11 motifs appear to regulate multiple r-protein genes (Table 1), so it is not possible to make as confident a prediction. It might seem most likely that the ligand would be encoded by the gene immediately downstream of the r-leader. However, this is a poor heuristic for predicting r-leader ligands; in fact, among r-leaders that regulate multi-gene operons in *Escherichia coli* or *Bacillus subtilis*, the ligand is most often not encoded by the immediately downstream gene (Additional file 1: Table S4) [7]. In early research, it was proposed that r-leader ligands are proteins that bind rRNA independently of other proteins [5]. Such primary-binding proteins have been distinguished from secondary-binding proteins, which bind rRNA only in the presence of other r-proteins [31, 32]. However, while primary-binding proteins are the most typical r-leader ligands, some subsequently validated r-leaders bind secondary-binding proteins, e.g., S2, S6:S18 and L25 [7, 31, 32].

We therefore exploited the observation that many r-leader ligands are common to multiple organisms, e.g., *E. coli* and *B. subtilis* [7] (Additional file 1: Table S4). We used this observation for motifs that regulate multiple genes where one of these genes encodes an r-protein that has been previously established as an r-leader’s ligand. For such motifs, we predict that the previously established r-protein is also the ligand of the new motif. To date, no exception to this assumption has been experimentally established. However, there is no guarantee that this situation will hold for all cases in the future (Additional file 1: Supplementary text). Despite this caveat, the assumption presents the best currently known method to predict r-leader ligands.

For 10 of the remaining 11 motifs that regulate more than one ribosomal gene, exactly one of the regulated genes has previously been identified as the ligand of an r-leader in a distinct bacterial lineage (Table 1, “Ligand basis” is “Prior”). One motif remains that is upstream of multiple genes, none of which encode a previously established r-leader ligand. Our best guess for this motif’s ligand is the immediately downstream gene, encoding L2 (Table 1).

It is important to emphasize that our predictions of ligands vary greatly in confidence, given the caveats mentioned in the preceding paragraphs. Because of the potential for incorrect ligand hypotheses, we list additional genes associated with the various motifs (Table 1 and Additional file 2). An additional point is that it is possible that some RNAs that regulate r-proteins in *cis* do not function by binding any r-protein [33].

Given a tentative ligand, we investigated possible mimicry of the rRNA by comparing the r-leader to the rRNA’s protein binding site (see Methods). Because atomic-resolution structures of our r-leaders are not available, we conducted this analysis using conserved features in the sequence and secondary structure. We were particularly interested in rRNA nucleotides involved in the binding interface that are highly conserved, as these nucleotides and their structural contexts are most likely to be adopted by an r-leader in order to imitate the rRNA. In performing this analysis, it is important to consider the possibility that apparent similarities between one or two conserved nucleotides might arise by chance. We conducted these analyses manually, because no model exists for quantitatively evaluating statistical significance. For r-leaders regulating multiple genes, we analyzed both the protein encoded by the immediately downstream gene and the most-likely protein ligand, if these are different. The consideration of multiple potential ligands and associated rRNA regions increases the chances of finding spurious similarities in local secondary structures, and therefore we did not attempt a comparison of more than two proteins. Evidence of possible mimicry is presented where the similarity between the r-leader and rRNA is striking (see Methods).

In the following text, we discuss specific findings about the 20 novel r-leaders. Additional details are in Additional file 1: Supplementary text.

### L31 r-leaders: five motifs for one r-protein

We found five motifs that likely bind the bacterial L31 r-protein (Table 1, Fig. 2), for which no r-leader has previously been identified. All five L31 motifs consist of a single hairpin, but the differing patterns of sequence conservation and bulge locations suggest that the motifs are structurally unrelated. However, it is conceivable that elucidation of the binding determinants or an atomic-resolution structure would reveal currently obscure similarities.

**Figure 2.**
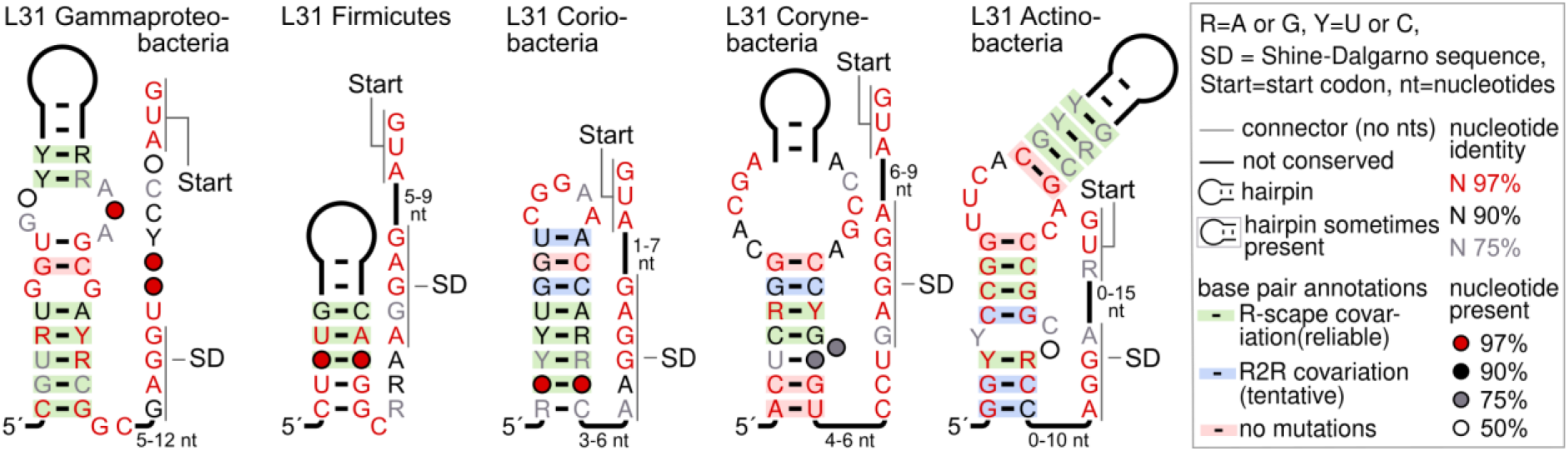
Conserved features of L31 r-leader motifs. The bacterial lineage is indicated. The inset explains annotations used throughout this paper. Information on our analysis of covariation is in Methods. Only R-scape annotations are statistically reliable.

The predicted L31 r-leaders span multiple phyla, although each individual r-leader motif is restricted to one phylum. We found novel L31 r-leaders in *E. coli* and *B. subtilis*, which is surprising, given the extensive study of these organisms.

The precise structure and role of the L31 protein in the ribosome has been unclear [34, 35]. A truncated form of L31 is likely a side effect of ribosome purification methods, and may have contributed to confusing data about L31 function [34]. Recent results suggest that L31 is part of a bridge between the small and large subunits and interacts with 5S rRNA, 16S rRNA and r-proteins L5, S13, S14 and S19 [35]. The ability of the 30S head domain to swivel is accommodated via changes in the flexible structure of L31 and its intermolecular interactions [35, 36].

Many bacteria have two L31 genes, where one of these two genes encodes a zinc-binding protein containing the amino acid sequence CXXC, where X is any amino acid [37]. These paralogous genes were proposed to contribute to regulation of zinc homeostasis [37, 38]. It is conceivable that the L31-associated motifs regulate L31 genes as part of zinc homeostasis, and are not L31-binding r-leaders. If this hypothesis is true, all five motifs would most likely have the same biochemical function, i.e., would either all regulate zinc-binding L31 proteins, or all regulate non-zinc-binding L31 proteins. However, this is not the case (Additional file 1: Supplementary text, Additional file 1: Table S5). Therefore, this hypothesis is probably incorrect, although we cannot rule out the possibility that some of the five L31 motifs have different biological functions from others. Curiously, we notice that most organisms contain zero or one L31 motif, for all five r-leader motifs (Additional file 2). Given that many organisms have two L31 genes that could be regulated, we are uncertain as to why only one is associated with an r-leader.

### Imitation of the rRNA: L20, eL15, S15 and S4 r-leaders

We found motifs that exhibit similarity to the rRNA’s binding site for either the L20, eL15, S15 or S4 r-proteins, and they are our best candidates for rRNA mimicry. These similarities are based on several conserved nucleotides in the rRNA binding site that resemble conserved r-leader nucleotides. Additional support derives in some cases from similarities between our r-leaders and previously published r-leaders whose rRNA mimicry was established in prior work.

R-leaders that bind L20 have been previously established in Gammaproteobacteria and Firmicutes [7]. These leaders each exhibit two conserved regions that correspond to two parts of the relevant binding site in the rRNA: two consecutive G-C base pairs and an AA dimer [7, 39] (Fig. 3a, Additional file 1: Figure S6). The two regions dock with each other in the ribosome (PDB model 4V85). We found a candidate L20 r-leader in Deltaproteobacteria that appears to conserve the same two regions of the rRNA binding site of L20, but in yet another structural scaffold (Fig. 3a). In the new motif’s AA dimer, one A nucleotide is predicted as pairing, but the existence of the base pair is unclear, as it is strictly conserved and therefore does not exhibit covariation. The two conserved elements are separated by a helix of several consecutive base pairs, which could bend to accommodate the docking interaction. Because of the apparent similarity to previously established L20 r-leaders that mimic the rRNA (Additional file 1: Figure S6), we suspect that the Deltaproteobacteria L20 motif also uses rRNA mimicry.

**Figure 3.**
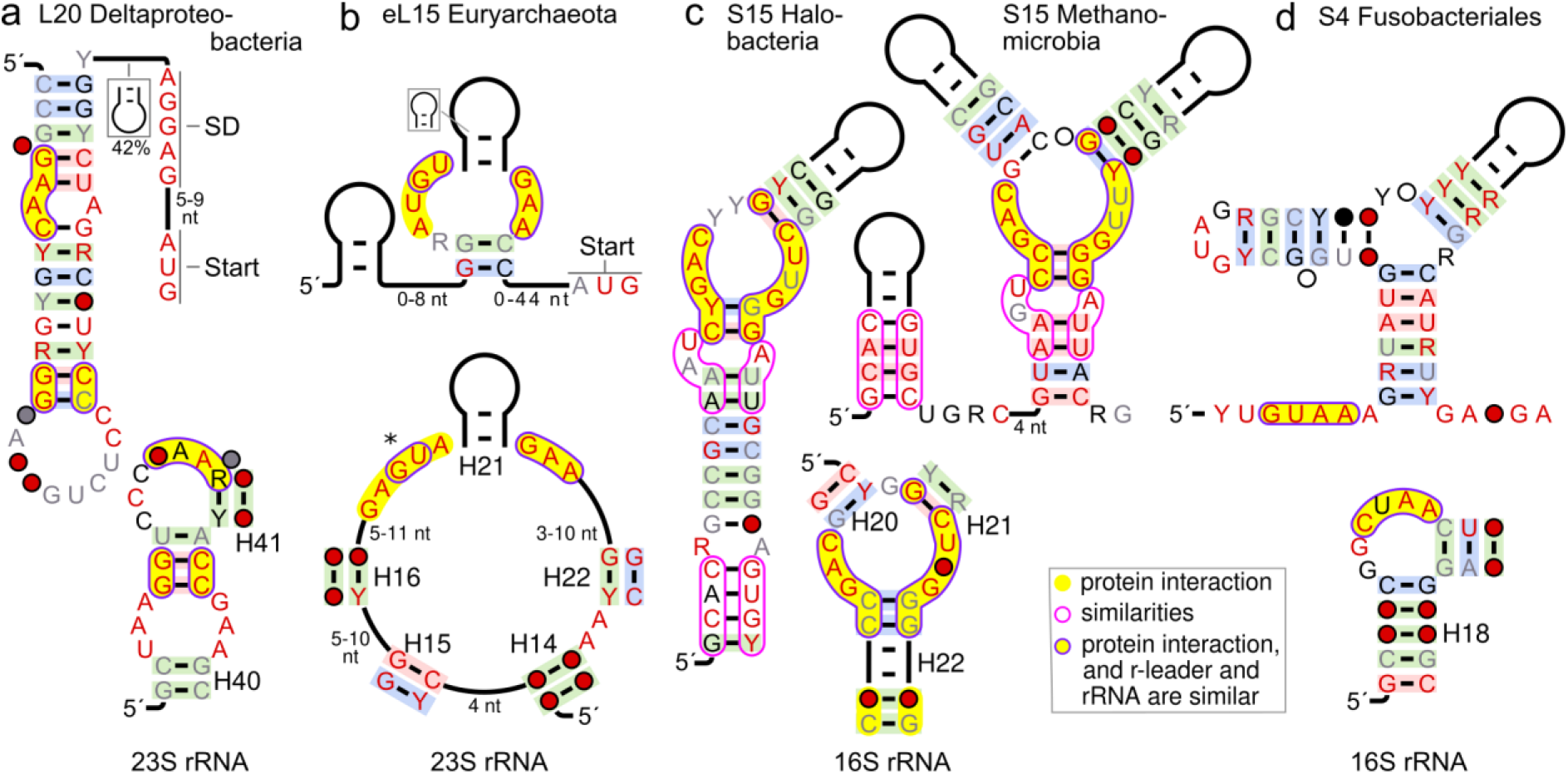
Proposed rRNA imitation by r-leaders. Upper row: novel r-leader motifs. Lower row: rRNA binding site for the relevant r-protein. Annotations: as in Fig. 2. Yellow shading: nucleotides that interact with the relevant r-protein. Purple outline with yellow shading: interacting nucleotides that are similar in r-leader and rRNA. Purple outline: similar r-leader nucleotides that do not resemble the rRNA. H20-H40: *Thermus thermophilus* helix numbers from [59] or [60]. (**a**) L20 motif. (**b**) Archaeal motif for the eL15 protein. The asterisk marks a region of the bulged-G module that is only partially similar in the r-leader (see text). (**c**) S15 motifs in Halobacteria and Methanomicrobia. (**d**) S4 motif from Fusobacteriales.

An r-leader motif that likely binds eL15 (the eukaryotic and archaeal L15 r-protein) (Fig. 3b) exhibits possible similarity to the eL15 binding site in the archaeal ribosome. The rRNA nucleotides that directly bind the eL15 protein (Fig. 3b) form a bulged-G module structure [40] (also called a Sarcin-Ricin loop or E-loop), a common structure in RNAs that is often associated with the binding of proteins [40]. The eL15 r-leader motif contains conserved nucleotides that are similar to a bulged-G module, but some nucleotides that are highly conserved in bulged-G modules are altered or missing in the r-leader motif (Fig. 3b, asterisk). Therefore, despite similar nucleotides, it is unclear whether this r-leader imitates the rRNA.

The strongest candidates for mimicry are two S15 r-leaders. The S15 protein is the target of four experimentally confirmed and two predicted bacterial r-leaders [41]. Experimental analysis of the four validated S15 r-leaders revealed how RNA-protein interactions can evolve over large evolutionary distances in which the r-leaders evolve distinct—yet partially related—strategies to bind the S15 protein [15]. Each of these motifs mimics one of two sites in the rRNA: (1) stacked G-C, G-U base pairs, or (2) conserved nucleotides within a three-stem junction [7] (Fig. 3c, Additional file 1: Figure S10).

We found two related archaeal S15 r-leaders (Fig. 3c). Although their secondary structures differ, a part of each motif closely resembles the multistem junction in the rRNA, strongly suggesting imitation (Fig. 3c). In addition to their mimicry of the rRNA, the motifs have regions that resemble each other, but do not resemble the rRNA (Fig. 3c, non-filled purple boundaries). One of these regions is a stem that occurs in different structural contexts in the two motifs. Given that the stem is conserved despite a structural rearrangement, it is likely that it is functionally important. However, the significance of the common regions to r-protein binding or gene regulation is unclear. Regardless, the two archaeal S15 motifs expand the scope of studying the myriad r-leaders for this r-protein to the domain Archaea.

An S4 r-leader that likely mimics the rRNA is described in the next section.

### S4 r-leaders

S4 is the experimentally confirmed target of known, structurally unrelated r-leaders in Gammaproteobacteria [42, 43] and Firmicutes [14, 44], although none are present in Clostridia, a class of Firmicutes [14]. The Gammaproteobacteria motif does not mimic the rRNA, but a conserved GUAA sequence in the Firmicutes motif was found to imitate the rRNA [14, 45].

Four of the novel r-leader motifs most likely bind the S4 r-protein (Table 1, Fig. 3d, Fig. 4a). We observed potential for the S4 motif in Fusobacteria to imitate the rRNA’s binding site (Fig. 3d). The conserved GUAA sequence in the previously published Firmicutes r-leader (Additional file 1: Figure S8) is shared by the Fusobacterial motif. Moreover, the GUAA sequence occurs in the context of other structural and sequence similarities between the two motifs, so this GUAA sequence in the Fusobacteria motif presumably also resembles the rRNA. A detailed comparison between the motifs, which also describes important differences between them, is in Additional file 1: Supplementary text.

**Figure 4.**
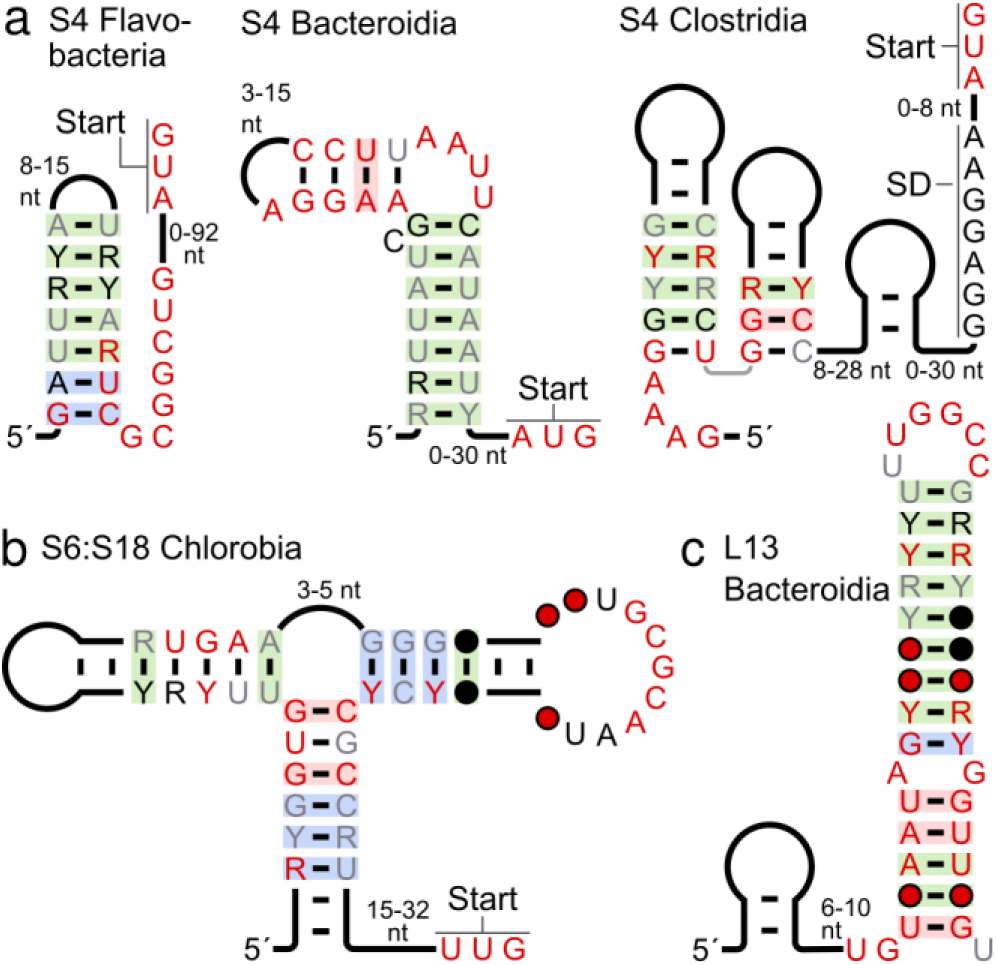
S4, S6:S18 and L13 r-leader motifs. (All 20 motifs are shown in Additional file 1: Figures S2-S11). Annotations are as in Fig. 2. (**a**) S4 motifs that do not appear to imitate the rRNA. (**b**) S6:S18 motif in Chlorobi. (**c**) L13 in Bacteroidetes.

The new S4 r-leader in Clostridia (Fig. 4a) has a conserved GAAA sequence (5′ end of molecule) that could resemble the aforementioned GUAA sequence, but this sequence is less similar to that of the rRNA. Therefore, it is unclear whether this r-leader might also imitate the rRNA. The two remaining S4 motifs, occurring in the related lineages Bacteroidia and Flavobacteria (Fig. 4a), exhibit no meaningful similarity to one another (other than the fact that they are both hairpins), to the previously published S4 r-leaders or to the rRNA.

### S6:S18 r-leader

The S6 and S18 proteins function as a dimer [23, 24]. A previously known r-leader binding S6:S18 (Additional file 1: Figure S9) is very widespread in bacteria [23, 24], but missing in certain lineages such as the phylum Chlorobi [24]. We found an r-leader (Table 1, Additional file 1: Figure S9) that most likely also binds S6:S18 and is restricted to the phylum Chlorobi (Fig. 4b). The widespread S6:S18 r-leader exhibits a clear similarity to the rRNA binding site (Additional file 1: Figure S9) and the relevant nucleotides are essential for binding the protein dimer [23, 24]. By contrast, the new Chlorobi version of this motif does not appear to resemble the rRNA.

### Archaeal r-leaders: S15, eL15, L4 r-leaders

Four archaeal r-leaders are among the novel motifs (Table 1). Three, which are expected to bind the S15 or eL15 proteins, were discussed above. We noticed some poly-U stretches immediately after S15 r-leaders in Methanomicrobia (Additional files 2 and 3) that could correspond to intrinsic termination signals, which are still not well understood in archaea [46, 47]. The fourth archaeal r-leader appears upstream of operons containing the L4 r-protein (Additional file 1: Figure S2, Additional file 1: Supplementary text), which is the ligand of a previously established r-leader in bacteria [7].

### Other r-leaders

We now note properties of the remaining candidate r-leaders beyond those summarized in Table 1. Some additional details are discussed in Additional file 1: Supplementary text. An experimentally verified L13 r-leader occurs in *E. coli* [7, 48], and a structurally distinct L13 r-leader motifs was previously predicted in Firmicutes [29]. We found a third, structurally unrelated L13 r-leader in Bacteroidetes (Fig. 4c).

A predicted L17 r-leader occurs in both Actinobacteria and Proteobacteria. This motif is the only candidate r-leader we found that is clearly used by more than one phylum, suggesting that such widespread r-leaders are unusual among still-undiscovered r-leaders. A previously predicted RNA element occurring in distinct organisms and called the “L17DE motif” [25] is consistently found downstream of L17 genes, and could bind the encoded proteins or those of another gene in the upstream operon. The L17DE motif is restricted to the phylum Firmicutes.

## Discussion

Using a comparative genomics approach, we found 20 r-leaders in multiple lineages of bacteria and archaea. These predictions are supported by covariation evidence that has proven reliable in past studies, and are in agreement with TSS positions determined by high-throughput experiments. Nonetheless, experimental study of our candidates will be worthwhile to validate the predictions. Moreover, experimental analysis could lead to a better understanding of their biology and implications for RNA-protein interactions.

The identification of 20 predicted r-leaders represents an increase of more than 50% over the 35 previously published r-leaders (Additional file 1: Table S1) and suggests that many more r-leaders remain undiscovered in biology. Thus, further application of comparative genomics or other methods would likely uncover additional r-leaders. Additionally, our discovery of several r-leaders for r-proteins that were not previously known as r-leader ligands (i.e., L2, eL15, L17, L31, S16) suggests that current knowledge of what r-proteins can function as r-leader ligands is incomplete.

We found only one motif that spans more than one phylum. This agrees with analysis of Gammaproteobacteria suggesting that r-leaders are typically not widespread [43]. Since many phyla remain understudied, our results suggest that a significant number of r-leaders likely remain undiscovered in these other phyla. However, the L17 motif in Alphaproteobacteria and Actinobacteria shows that there is still room for the discovery of additional widespread r-leaders.

The four r-leader motifs in Archaea represent a significant increase in the number of known *cis*-regulatory RNAs in these organisms. Archaea share many characteristics of both eukaryotes and bacteria [47]. The novel r-leaders present rare opportunities to learn about how *cis*-regulation works at the RNA level in Archaea, and, by extension, how transcription and translation processes may be co-opted for use in regulation.

The hypothesis that r-leaders will mimic rRNAs was presented soon after their discovery [5]. However, while many r-leaders do indeed imitate the rRNA binding site, several do not [7]. In the absence of rRNA imitation, distinct amino acids in the relevant r-proteins might be conserved that are not important for the r-protein’s primary function. In this context, rRNA imitation might be an economical strategy that reduces the need for sequence conservation.

Of the 20 novel r-leader motifs, only 25% exhibit plausible similarities to the rRNA (Table 1, Fig. 3). By comparison, of 20 previously analyzed r-leaders [7], 50% show good evidence of rRNA imitation [7]. This comparison could suggest that mimicry is, in fact, unusual in r-leaders, or that it is only common in widespread r-leader motifs, whereas the new r-leader motifs are generally restricted to one phylum. Another observation is that r-leaders of some proteins seem to consistently copy the rRNA, e.g., S15 and L20. Other proteins are inconsistent, e.g., the S6:S18 dimer has one previously published motif [23, 24] with clear similarity, and another (our Chlorobi motif) with no apparent similarity. Perhaps some protein-binding sites are more conducive to imitation than others.

We note that there are technical reasons that could account for our inability to establish a convincing similarity between r-leader and rRNA in some cases. First, comparisons between r-leaders and rRNA are more difficult because of uncertainty about the ligand and the absence of atomic-resolution structural information for the r-leaders. Second, in some cases, the rRNA binding sites show only a limited number of conserved nucleotides, or are not well conserved. In such cases, it is easy to find positions in the r-leader that might be similar, but difficult to establish that the similarity is meaningful. Therefore, it is possible that additional research will reveal more cases of structural mimicry among our r-leaders.

Previous work has already shown that there are r-proteins whose r-leaders exhibit multiple, distinct primary and secondary structures. Examples in *E. coli* and *B. subtilis* are the r-leaders for L20, S4 and S15 [7] (Additional file 1: Table S4). Considering all bacteria, four distinct r-leaders binding S15 were previously known [41]. Our work adds an archaeal version of S15, with two sub-types, and potential r-leaders for S4, L4, L13, L20, S4 and S6:S18 that add r-leaders to previously established protein ligands. Additionally, there are now five motifs associated with L31 genes. This plethora of solutions for binding a consistent protein could suggest that such interactions are easy to evolve, and creates fertile input for studies on the evolution of protein-RNA interactions [15].

## Conclusions

This study presents 20 novel r-leaders in bacteria and archaea. The predictions are supported by covariation evidence, gene associations, and, in many cases, by high-throughput TSS experiments, and are thus a good starting point for detailed experimental investigation. With an increase of more than 50% in the number of known or predicted r-leaders, these results suggest that many more r-leaders remain undiscovered in bacteria and archaea.

R-leaders offer valuable opportunities to better understand RNA-protein interactions. The newly found r-leaders include multiple RNA structures that likely share a common ligand, or whose ligand is also the ligand of a previously established r-leader. Thus, the r-leaders represent alternate evolutionary solutions to the same protein-recognition problem. Additionally, some of the r-leaders exhibit a similar structure to that of the rRNA binding site, although for many r-leader motifs, no compelling similarity is apparent. The new r-leaders thus represent a foundation for different types of studies on r-leaders, gene regulation, ribosome assembly and RNA-protein interaction.

## Methods

### Databases and software

We used sequence data from the RefSeq nucleotide database version 72 [49] and metagenomic data predominantly from IMG/M [50] and GenBank [51]. Genes were annotated as previously described [26]. Known RNAs were annotated using the Rfam Database [52] version 14.0 and papers on r-leaders [7, 14, 23, 29, 41, 43, 48]. Homology searches were conducted using Infernal version 1.1 [53], and alignments were analyzed for additional secondary structure using CMfinder version 0.4 [26, 54] and R-scape [55]. RNAs were drawn using R2R [56] and Inkscape [57].

### Covariation

Several existing algorithms assess covariation in an alignment. Unfortunately, currently available computational approaches cannot consistently handle problems such as incorrect alignments, which can create misleading, invalid covariation. We therefore ultimately evaluated covariation manually. However, we used R-scape [55] with the -s option as a guide. Assuming a valid alignment, R-scape is a statistically well-founded indicator of covariation. R-scape measures the statistical significance of a covariation signal using a random model of evolution that accounts for phylogenetic signals that can confound covariation analysis. Base pairs exhibiting statistically significant covariation are then reported. All 20 r-leader motifs exhibited statistically significant covariation according to R-scape (Additional file 1: Table S2). An issue with R-scape is that it does not consider small covariation signals in multiple base pairs that together could provide compelling evidence of covariation. So, for the diagrams, we also depicted R2R’s [56] simplistic and more permissive method to detect potential covariation. R2R reports covariation when there are at least two Watson-Crick or G-U base pairs among the sequences that differ at both positions and fewer than 10% of the sequences have non-canonical base pairs (i.e., base pairs that are neither Watson-Crick nor G-U) at those positions. Importantly, R2R’s annotations are neither reliable nor statistically well-founded [56], and were therefore not used to draw conclusions about candidates. To allow manual or other analyses of our predicted r-leaders, we provide our alignments (see Data Availability). Based on our manual analysis, we have not drawn stems we deem dubious, and note a few hairpins that are uncertain (Additional file 1: Supplementary text).

### Comparison of structures in rRNAs and r-leaders

As part of the analysis of whether the new r-leaders structurally imitate rRNA, we used rRNA alignments available in the Rfam Database [52] version 14.0 (Additional file 1: Table S6). To more easily connect nucleotides in ribosomal crystal structures to alignment columns, the alignments were modified by adding rRNAs from the relevant crystal structure in Protein Databank (PDB) [58] using Infernal. Information on rRNA nucleotides that directly interact with a given r-protein were extracted from relevant studies (Additional file 1: Table S7) and inferred with the PyMOL (Schrödinger, LLC) version 2.0 function “Action: find > polar contacts > to other atoms in object”. We used a previously established crystal structure of the ribosome (PDB accession: 6GZQ) to conduct this analysis. For the eL15 r-leader, we used an archaeal ribosome (PDB accession: 1S72). Structural comparisons were performed manually based on drawings of conserved sequence and secondary structure features.

### Comparison of r-leader motifs

We compared r-leader motifs with previously established r-leaders and with each other, in order to determine if they were novel. As with comparisons to rRNA, we conducted these comparisons manually based on the conserved primary and secondary structural features of the relevant alignments. We paid particular attention to r-leaders that potentially had the same r-protein ligand, based on proteins encoded by regulated genes, as these r-leaders would be most likely to be similar.

### Operon analysis

Information on regulated operons in Table 1 were estimated manually. Due to multiple factors, it is not straightforward to analyze this information automatically. Such complications include the high number of truncated contigs in metagenomic and shotgun sequences, errors in automated gene analysis (esp. the annotation of spurious genes) and natural variation in the distance between genes in operons, for example. Therefore, we initially assumed that co-transcribed genes can be up to 500 nucleotides apart, and considered only r-leader homologs where at least 8 Kb of sequence was available downstream of the RNA. We did not include genes that are consistently located far away from the previous gene, inconsistently positioned or often absent. Such additional genes that might be regulated are available in Additional File 2.

## Ethics approval and consent to participate

Not applicable.

## Consent for publication

Not applicable.

## Availability of data and materials

All data generated or analyzed during this study are included in this published article and its supplementary information files, or referenced therein. R-leader motif alignments are available (in addition to Additional files 2-4) in an Rfam-Database-like [52] format at https://bitbucket.org/zashaw/zashaweinbergdata/src/master/r-leader/, which is part of the repository at https://bitbucket.org/zashaw/zashaweinbergdata/src/master/.

## Authors’ contributions

ZW conceived of and supervised the study. IE conducted the study. Both authors wrote and approved the final manuscript.

## Acknowledgements

We are grateful for computer time provided by the Center for Information Services and High-Performance Computing (ZIH) at TU Dresden. We also thank Christina E. Weinberg for a critical reading of the manuscript, and Michelle Meyer and bioinformaticians at Leipzig University for helpful comments.

## Additional files

### Additional file 1

Supplementary text, tables and figures. (PDF 2,009 KB)

### Additional file 2

Alignments and information on downstream genes and taxonomy for all predicted r-leader motifs. (PDF 7,447 KB)

### Additional file 3

Archive of machine-readable alignments for all r-leader motifs that include flanking sequences and metadata. The alignments are stored in Stockholm format, which are text files that can be opened in any text editor (using a fixed-width font) and interpreted by software packages such as Infernal [53]. (ZIP 1,367 KB)

### Additional file 4

Archive of machine-readable alignments for all r-leader motifs that include only nucleotides within the predicted motifs, and do not include any metadata. The alignments are appropriate for tasks such as performing homology searches or re-analysis of our predicted secondary structures. The alignments are stored in Stockholm format. (ZIP 323 KB)

