## Additional file 1 for "Discovery of 20 novel ribosomal leaders in bacteria and archaea"

### Additional file 1: Supplementary text, tables and figures

### For:

#### Supplementary Text

##### Predicting the ligands of new r-leaders based on the ligands of previously established r-leader

As noted in the main text, we wish to generate a hypothesis for the ligand of each of the 20 newly found r-leaders. For r-leaders that are predicted to only regulate genes encoding one r-protein, it is clear that this r-protein is the most likely ligand. For r-leaders that regulate genes encoding multiple r-proteins, we proposed in the main text that the most likely ligand is whichever r-protein has previously been established as the ligand of an earlier r-leader. There is no experimentally established counter-example to this rule.

However, it is clear, given the relatively low number of r-leaders with validated ligands, that we cannot rule out the possibility that exceptions might be proven in the future. Also, whereas an *E. coli* r-leader exists whose ligand is L4, experiments suggested L4 does not bind the putative r-leader in *B. subtilis* that regulates similar genes [1]. While these results are suggestive, they do not establish that the ligand is not L4 [1]. First the experiments merely failed to show L4 binding, and did not validate a different ligand. Second, the experiments were conducted in a surrogate host. Despite these caveats, we believe that the use of previously established r-leader ligands is likely to provide good hypotheses for the ligands of newly predicted r-leaders.

##### Searching for differential RNA-seq (dRNA-seq) datasets

To search for relevant dRNA-seq datasets, we enumerated genera in which many examples of each of the 20 novel r-leader motifs occur. We then searched for dRNA-seq data in these genera using multiple strategies. The most successful strategy was to use Google Scholar [2] to search for the names of the various genera containing the r-leader motifs within papers citing the study that introduced dRNA-seq [3]. We also performed *ad hoc* Google Scholar searches. Our remaining strategy was to search for RNA-seq studies within the Short Read Archive (SRA) [4] on the NCBI Web site, especially for the term “dRNA-seq”. However, we found that the type of experiment was not consistently stated in the SRA metadata.

#### Additional notes on motifs, including motifs not described in the main text

### L2

We found an r-leader upstream of genes that encode L2 in Alphaproteobacteria (Figure S2). In *E. coli*, the L2 and L4 genes form an operon, and the L4 protein is the ligand of an r-leader. However, the Alphaproteobacterial operons lack the L4 gene. Therefore, we presume that the most likely ligand is that encoded by the immediately downstream genes, i.e., L2.

We regard this motif as a borderline prediction (Table 1), because of the relative lack of conserved nucleotides around the stems and relatively frequent non-conserved insertions (Additional files 2-4). Thus, although this motif likely functions as an r-leader, especially given the precedent that many bacterial ribosomal genes are regulated by r-leaders, the data are not as convincing as with other motifs.

### L4

The L4-Archaeoglobi r-leader motif has many examples that overlap protein-coding genes, which is unusual for *cis*-regulatory RNAs. Our conclusion is that the motif is very likely to correspond to an RNA with a conserved secondary structure, and that the motif remains a credible r-leader candidate. However, in view of the unusual locations of these r-leaders, we have marked the motif's assignment as an r-leader as relatively borderline (Table 1). We explain our reasoning in detail in the following text.

Roughly 54% of L4-Archaeglobi r-leader examples (254 sequences) are found within the coding regions of genes predicted to encode RNA methyltransferases defined by Pfam [5] entry PF02598. The remaining 46% of L4-Archaeglobi r-leader examples (215 sequences) do not occur inside such genes, or in other types of genes. We believe many of the 254 r-leaders do, in fact, overlap a coding region, i.e., this overlap is not the result of a genome annotation error. This conclusion follows because of the large number of overlapping predictions, and since we observed insertions in multiples of three nucleotides among affected r-leader sequences. Such insertions suggest a coding function because each codon is three nucleotides. Additionally, of the 254 sequences that overlap these genes, there are zero stop codons in the third reading frame (i.e., starting with the third nucleotide in each sequence, which corresponds to the reading frame of the methyltransferase genes). Based on the number of codons and the frequencies of the four nucleotides, the expected number of stop codons in random nucleotides would be 217.

It also appears that many L4-Archaeoglobi motif examples do not overlap the gene, i.e., the absence of the gene is also not a genome annotation error. For example, we chose a non-overlapping motif example (in RefSeq accession NZ_CM001555.1) and compared it and its surrounding genomic nucleotides to a protein encoding an overlapping gene (protein accession WP_049993178.1) using BLASTX (which internally translates nucleotide sequences into amino acid sequences in all frames). We did not find any similarity with an E-value better than 1. This result provides additional evidence that this sequence does not code for an RNA methyltransferase like Pfam entry PF02598.

We also analyzed whether any of the 215 sequences that do not overlap a gene contained stop codons. Such stop codons would suggest the absence of coding function. We investigated all three reading frames on the sense strand, and found they all had many stop codons. In the first reading frame, there were 6,141 codons among the 215 sequences, of which 117 were stop codons. Based on the number of codons and frequencies of the four nucleotides, 206 stop codons would be expected. In the second reading frame, 313 of 6,104 codons were stop codons, versus an expected 212. In the third reading frame, 79 of 5,971 codons were stop codons, versus 200 expected. Overall, there is no clear evidence of coding function in these sequences.

One possible concern with this r-leader motif is that, in performing homology searches, we inadvertently included sequences that are not homologous. In particular, perhaps the L4-Archaeoglobi motif sequences that overlap the methyltransferase genes are not actually homologous to the motif sequences that do not overlap these genes. We therefore compared all overlapping sequences to all non-overlapping sequences using the blastn program from NCBI's BLAST [6] package with a word size of 7 and an E-value cutoff of $10^{-5}$. Out of the 254 gene-overlapping sequences, 252 had a match to a non-overlapping sequence that BLAST could find. In the reverse direction, BLAST was able to match 75 L4-Archaeoglobi motifs examples that do not overlap a methyltransferase gene against motif examples that do overlap genes. Since BLAST is not designed for RNA and does not consider secondary structure, it is noteworthy that it is able to detect so many similarities. We also used the Infernal software [7] to train a covariance model on the motif examples that overlap genes, and used it to search the sequences of non-overlapping motif examples. We found 175 (out of 215) of the sequences with an E-value better than $10^{-5}$, and 190 with an E-value better than $10^{-2}$. Therefore, there is strong evidence that the motif examples that overlap genes are homologous to the motif examples that do not overlap genes.

Additionally, we found that an alignment of the 254 sequences that overlap genes exhibits covariation according to R-scape—including E-values better than $10^{-5}$. An alignment of the 215 sequences that do not overlap RNA methyltransferase genes similarly exhibits covariation, with R-scape E-values below $10^{-5}$. Thus, each group of motif examples independently has evidence of covariation.

Thus, the L4-Archaeoglobi motif likely represents an RNA that sometimes occurs within a protein-coding gene, and sometimes does not. The RNA could therefore regulate the downstream gene from transcripts that include the overlapping methyltransferase genes. Since 46% of L4-Archaeoglobi RNAs neither overlap nor are near to methyltransferase genes, it seems unlikely that the function of the RNA is primarily related to these protein-coding genes. However, the unusual arrangements of this motif reduce our confidence L4-Archaeoglobi RNAs function as r-leaders (as reflected in Table 1).

Another possibility to consider is that the motif does not function as a *cis*-regulatory RNA and does not have a conserved RNA secondary structure. However, the covariation information is significant both quantitatively (Table S2 and above text) and qualitatively. Therefore we believe that the motif likely does function as an RNA.

## L13

This motif consists of two adjacent hairpins: a 5´ and a 3´ hairpin (Figure S3). Although the 3´ hairpin in this motif is well supported by covariation, the 5´ hairpin is less clear, due to a relatively high variation in lengths of unpaired regions, and the relative absence of strong sequence conservation. This combination of features allows the possibility that the base pairs arise by chance. In this view (that the base pairs are not biological, but rather arise by chance), the variable lengths of unpaired regions give the computer great flexibility in finding spurious base pairs located somewhere in a relatively long sequence. Thus, such variable lengths can be associated with false-positive stems. Arguing against this view, however, we do notice that even sequences that share the same lengths exhibit covariation within the putative 5´ hairpin. Since the sequences have similar lengths, they appear to be related, and the covariation within these sequences is less likely to arise by chance. Moreover, if the 5´ hairpin truly occurs in these related sequences, then it is reasonable to assume that it occurs in other sequences. Therefore, this 5´ hairpin is likely of biological importance to the host, although our level of confidence is not as high as with other hairpins in the drawings of the various motifs.

## L17

This motif (Figure S5) has a 5´ hairpin whose biological significance is not completely clear because of variation in lengths of unpaired regions and a relative lack of sequence conservation, similar to the case with the L13 r-leader motif, above. As with the L13 hairpin, the 5´ hairpin in the L17 r-leader motif exhibits covariation even among similar sequences. There are, however, two identical sequences in which no compelling stem can be found. We conclude that the hairpin in the L17 motif is likely to be biological, at least in most members of the motif.

## L20

The r-leader is immediately upstream of the L20 gene and immediately downstream of the L35 gene. In *E. coli* and *B. subtilis*, the L35 gene occurs immediately upstream of the L20 gene, but in these organisms, the relevant r-leaders occur immediately upstream of the L35 gene. Thus, the order of the genes is the same, but the r-leader in Deltaproteobacteria occurs between the genes. Given the precedents that L20 is an r-leader ligand, we hypothesize that the new Deltaproteobacteria motif also binds L20, and that the position of the L35 gene is therefore of little importance.

## L31

##### *Zinc proteins and L31 motifs*

To determine if the L31 motifs are likely to have a function related to the zinc binding properties of some L31 proteins, we analyzed the regulated proteins to determine how often the motifs regulated proteins that contained or did not contain the zinc-binding peptide motif. For the L31-Coriobacteria and L31-

Firmicutes motifs, 60% and 67% of the regulated genes encode proteins with the zinc-binding element (Table S5). Thus, there is a mixture of zinc-binding and non-binding proteins encoded by the genes that are apparently regulated by these motifs. Thus, these two motifs probably have no relationship to zinc levels.

In contrast, the other three L31 motifs are consistently associated with one type of protein (Table S5), but the association is not consistent across these three motifs. In particular, while the motifs in Gammaproteobacteria and Actinobacteria are almost always associated with proteins that appear capable of binding zinc, the Corynebacteriacaea L31 motif only regulates genes encoding proteins that lack the zinc-binding pattern. It is possible that these three L31 motifs do act in zinc regulation, but it seems mostly likely that all five L31 motifs should have the same biochemical function. Moreover, we note that there is no organism that contains more than one type of these motifs. Therefore, it does not appear to be the case that one motif regulates zinc-binding proteins, while another motif regulates proteins that do not bind zinc. Thus, if it is indeed the case that all five motifs have the same biochemical function, these data suggest that the L31 motifs are not likely to play a role in zinc homeostasis.

##### *Other information on L31 r-leaders*

Three motifs are found in the phylum Actinobacteria, specifically those distributed in the taxa Actinobacteria, Coriobacteria and Corynebacteriaceae (Table 1, “Lineage”). However, these three motifs occur in distinct organisms; no single organism contains more than one type of L31 motif.

Among the L31 motif examples in Firmicutes, the sequence CUU is found often immediately upstream of the conserved hairpin. This sequence could bind the downstream gene’s Shine-Dalgarno sequence. However, a much larger number of motif instances lack the conserved CUU. Therefore, its significance is unclear. We have depicted a subset of L31 r-leaders in Firmicutes that have a CUU extension on their 5´ ends (Figure S7).

There is a potential stem in the L31-Gammaproteobacteria motif involving the SD sequence. The 5´ side of the stem involves nucleotides in the region depicted as “5-12 nt” (Fig. 2) and the GC dimer that occurs immediately 5´ to that region. The stem’s 3´ side involves nucleotides in the SD sequence and the U nucleotide immediately 3´ it. We did not indicate this stem in drawings or alignments, because it is not supported by covariation. However, it is plausible and might form part of a regulatory mechanism.

The L31-Gammaproteobacteria motif has the sequence UUCUGUAU(5-7nucs)GCC 40-120 nucleotides upstream of many r-leaders. However, these sequences seem to be associated with upstream genes that encode a DEAD-like helicase. When these genes are absent, the sequence is not found. Therefore, we do not believe that this partially conserved sequence is related to the L31-Gammaproteobacteria r-leader motif.

## S4

Although Firmicutes have only one previously established S4 r-leader, two distinct multiple-sequence alignments have been prepared of this r-leader (Figure S8). The first alignment to be published [8] was based on a comparative analysis of sequences upstream of S13 genes. A subsequent study [9] considered experimental data [10, 11] and adjusted the earlier alignment to conform to these experimental results. Both of these alignments exhibit a highly conserved GUAA sequence nearby to, and 5´ to a hairpin. In both alignments, there is a conserved GCU in the hairpin's terminal loop. In the earlier study [8], there is also a conserved YGA sequence 3´ to the hairpin, which is not aligned in the second study [9].

Our S4 motif in Fusobacteria has the GUAA sequence 5´ to a stem that contains a three-stem junction. (The motif has additional conserved nucleotides 5´ to the GUAA sequence, but this region is not conserved in the Firmicutes r-leader.) 3´ to the enclosing stem in the new Fusobacteria motif is a conserved YGA sequence, resembling the Firmicutes motif in the earlier study's [8] analysis. Oddly, the GUAA and YGA sequences are shifted in their positions relative to the stem when comparing the Firmicutes motif to the Fusobacterial motif. This dissimilarity cannot be resolved by removing or adding base pairs. The Fusobacterial motif does not have a GCU sequence in either of the terminal loops that the stem encloses. However, one of the stems does have a conserved GUA sequence. It is not clear if these conserved sequences are related.

The four novel motifs differ in terms of which genes they regulate in addition to the S4 gene (*rpsD*), but the regulated genes for each new r-leader are similar to those of some previously published r-leaders. Two S4 r-leaders have been published. The Gammaproteobacteria S4-binding r-leader occurs immediately upstream of the S13 gene (*rpsM*), and S4 is encoded by the third gene in the regulated operon [12], with a gene encoding S11 in between. The relevant r-leader in Firmicutes is immediately upstream of the S4 gene, and the S13 and S11 genes are located elsewhere in the genome. Among the novel r-leaders, the Fusobacteria and Clostridia S4 motifs (Table 1) occur upstream of the S13 gene, with S11 and S4 genes located further downstream. These r-leader motifs are also immediately upstream of the L36 gene. These characteristics of the Fusobacteria and Clostridia S4 motifs are similar to those of the Gammaproteobacterial r-leader. Two other new r-leader motifs (in Bacteroidia and Flavobacteria; see Table 1) are found directly upstream of the S4 gene (*rpsD*), and genes encoding S13 and S11 are found immediately upstream of the r-leaders. This gene order is the same as in *E. coli,* but in that organism, the r-leader is upstream of the S13 gene.

## S6:S18

The start codon downstream of S6:S18-Chlorobi r-leaders is conserved as UUG in annotated positions (Figure S9). Although gene annotations often contain incorrect start codons, the consistency of the predictions in all sequences suggests that the start codon was correctly annotated.

## S15

The new motif upstream of S15 genes in Flavobacteria (Figure S10) has strong evidence to support its assignment as an RNA, but its function as an r-leader is less clear. The sequences belonging to the S15-

associated motif in Flavobacteria contain a non-essential hairpin that exhibits covariation. The typical sequence pattern (TAxxTTTG) corresponding to a Flavobacteria promoter [13] occurs immediately upstream of the motif. The hairpin follows in most sequences, but many sequences have a short sequence (usually fewer than 6 nucleotides) that does not form base pairs. 3´ to the hairpin region are several conserved nucleotides that end in the start codon. Given the promoter sequence and the covariation in the hairpin, it is clear that this motif is transcribed as RNA. However, sequences missing the hairpin thereby lack any demonstrable secondary structure, while all r-leaders studied so far are believed to exhibit a conserved secondary structure in all cases. Moreover, we do not find any compelling similarity between the conserved nucleotides and the rRNA binding site for S15. Therefore, it is unclear whether this motif functions as an r-leader.

There are some positions within the archaeal motifs that could correspond to stacked G-C, G-U base pairs, one of the conserved features of the rRNA's S15 protein binding site. However, we judged the similarity to be uncompelling, given that this is such a simple pattern.

## S16

The S16 motif in Flavobacteria (Figure S11) is consistently located upstream of genes encoding S16. Downstream of the S16 gene are usually *rimM* genes. *rimM* genes encode proteins involved in 16S rRNA processing, and do not encode an r-protein. Despite the presence of *rimM* genes, this motif likely functions as an S16-binding r-leader for two reasons. First, it is less clear that the *rimM* genes are part of the regulated operon, due to an inconsistent distance between the genes (often close to 300 nucleotides) and occasional cases in which the *rimM* gene is missing. If the *rimM* genes are, in fact, not part of the operon, then it is clear that they have no relevance for the function of the RNA motif.

Second, previously validated r-leaders are known to regulate genes that do not encode r-proteins [12]. For example, the experimentally validated L20 r-leader in *B. subtilis* is found immediately upstream of *infC* genes, which encode translation initiation factor 3 [12]. However, this r-leader binds the L20 r-protein, which is actually encoded by the third gene in the operon. Another similar example is that of the S6:S18 r-leaders validated in *E. coli* and *B.* subtilis [12]. This r-leader binds S6:S18, but regulates genes that encode single-strand-DNA-binding proteins. Thus, there are at least two established r-leaders that bind r-proteins, but regulate genes that do not encode r-proteins.

Thus, the *rimM* genes might not be part of the operon regulated by the S16-associated r-leaders. Moreover, if the *rimM* genes are part of the operon, this would not contradict the hypothesis that the motif is an S16 r-leader, given the precedents mentioned in the previous paragraph for L20 and S6:S18 r-leaders.

The S16 r-protein is not established or proposed as the ligand of any previously published r-leader.

#### Supplementary Tables and Figures

**Table S1.** Previously published or predicted r-leaders. This table is expanded from a previously published review paper [12], using data from Rfam [14] and a recent paper [15]. The table includes only motifs that are either experimentally confirmed or have a published predicted multiple-sequence alignment. “R-protein ligand”: the experimentally verified or proposed ligand protein, according to the original paper. Multiple r-leaders that exhibit distinct structures and bind the same protein ligand are listed in separate rows. In some cases, the ligand has not been proposed. In such cases, the products of the proximal genes in the downstream operon are listed. Information on experimentally confirmed r-leaders is available in a previous review [12]. “Lineage”: the taxon that contains known examples of the r-leader. “Alignment citation”: citations reflect works that produced multiple-sequence alignments including multiple homologs. If such an alignment is not available, an earlier paper describing an r-leader in a single organism is given. “Rfam Database accession” : accessions are given for r-leaders present in the Rfam Database [14]. Special annotations: (a) Although the L7Ae protein is part of the archaeal ribosome, it is also present in many other RNA-protein complexes [16, 17], so it is at least not purely an r-protein. Thus, the assignment of this RNA as an r-leader depends on the precise definition of the term “r-leader”. We included it in the table, since the published r-leader has been shown to regulate the L7Ae gene by binding L7Ae. (b) The alignment in the Rfam Database is independent of the alignment present in the cited work. The alignments might complement each other. (c) The motif occurs downstream of the associated genes, and therefore likely functions in the 3´ UTR. This motif is therefore technically not an r-leader, even though it might bind the L17 r-protein and participate in feedback regulation. (d) No alignment is available for this r-leader, which is known only in one organism. (e) These alignments exhibit little or no covariation, except in a Rho-independent terminator, so the true secondary structure is unclear, and it is difficult to compare them to the other predicted S7 or S12 r-leaders. Therefore, it is uncertain if these are truly distinct structures. (f) S10 refers to the product of the first gene in an operon that contains many genes that each might encode the ligand of this gene. The correct ligand has not been experimentally determined.

| R-protein ligand (confirmed or predicted) | Lineage | Alignment citation | Rfam Database accession (if any) |
|---|---|---|---|
| L1 | Archaea & Bacteria | [18] | |
| L4 | Gammaproteobacteria | [18] | |
| L7Ae (a) | Archaea | [16, 17] | |
| L10 | Bacteria | [18] | RF00557 (b) |
| L13, S9 | Firmicutes | [8] | RF00555 |
| L13 | Gammaproteobacteria | [15] | |
| L17 (c) | Firmicutes | [19] | RF01708 |
| L19 | Firmicutes | [8] | RF00556 |

| | | | |
|---|---|---|---|
| L20 | Firmicutes | [9] | RF00558 |
| L20 | Gammaproteobacteria | [18] | |
| L20 | *E. coli* (d) | [20] | |
| L21, L27 | Firmicutes | [8] | RF00559 |
| L25 | Enterobacteria | [21] | |
| L28 | Gammaproteobacteria | [15] | |
| L34 | Gammaproteobacteria | [15] | |
| S1 | Cyanobacteria | [22] | |
| S1 | Gammaproteobacteria | [18] | |
| S2 | *Pelagibacter* | [23] | RF01815 |
| S2 | Bacteria | [18] | RF00127 (b) |
| S4 | Gammaproteobacteria | [18] | RF00140 |
| S4 | Firmicutes | [9] | |
| S6:S18 | Bacteria (but not Chlorobi) | [24, 25] | |
| S7 | Gammaproteobacteria | [18] | |
| S7, S12 (e) | *Pelagibacter* | [23] | RF01823 |
| S7, S12 (e) | *Pseudomonas* | [26] | RF01773 |
| S7, S12 (e) | *Rickettsia* | [26] | RF01774 |
| S8 | Gammaproteobacteria | [18] | |
| S10 (f) | Firmicutes | [8] | |
| S15 | Actinobacteria | [27] | |
| S15 | Alphaproteobacteria | [27] | |
| S15 | *Chlamydia* (d) | [27] | |
| S15 | Firmicutes | [9] | |
| S15 | Gammaproteobacteria | [18] | RF00114 |
| S15 | *Thermus thermophilus* (d) | [28] | |
| S20 | *E. coli* (d) | [29] | |

**Table S2**. Base pair and covariation statistics of predicted r-leaders, including statistical support of RNA covariation from R-scape. Note: due to issues described in Methods, we additionally evaluated covariation information manually. This table provides the following information for each predicted r-leader (including the three r-leaders we found that strongly resemble previously established leaders). “Number of seqs.”: the number of instances of the r-leaders within the sequence databases we searched. Note: the 29,730 r-leader examples mentioned in the main text do not include the three motifs (L19-Flavobacteria, L25-Gammaproteobacteria and S10-Clostridia) that we concluded are very similar in conserved features to previously published r-leader motifs. “Avg. len.”: the average length in nucleotides of the motif. “All”: the total number of base pairs in the alignment. Note: this includes aligned columns that have gaps in the vast majority of motif members, since there is often significant variation in the lengths of stems. “Invariant”: the Watson-Crick or G-U base pair is not observed to change. (Corresponds to red shading in our diagrams.) “Covary (R2R)”: covariation is predicted by R2R, but not by R-scape. (Corresponds to blue shading in our diagrams.) “Covary (R-scape)”: covariation is predicted by R-scape (and possibly also by R2R). (Corresponds to green shading in our diagrams.) Covariation predicted by R-scape [30] is statistically significant and has an E-value < 0.05. (E-values are similar to p-values.) Note: base pairs in gappy columns are generally not drawn in the diagrams, but are reflected in the numbers in this table. Therefore, these numbers will generally be higher than the number of base pairs in the corresponding diagrams.

| r-leader | Number of seqs. | Avg. len. | Number of base pairs | | | |
|---|---|---|---|---|---|---|
| | | | All | Invariant | Covary (R2R) | Covary (R-scape) |
| L2-Alphaproteobacteria | 808 | 70 | 42 | 9 | 3 | 13 |
| L4-Archaeoglobi | 654 | 94 | 49 | 5 | 10 | 24 |
| L13-Bacteroidia | 1380 | 76 | 26 | 3 | 5 | 16 |
| eL15-Euryarchaeota | 1001 | 57 | 153 | 46 | 27 | 11 |
| L17-Actino-Proteobacteria | 548 | 56 | 26 | 6 | 6 | 13 |
| L19-Flavobacteria | 2361 | 28 | 9 | 3 | 1 | 5 |
| L20-Deltaproteobacteria | 202 | 68 | 25 | 4 | 12 | 6 |
| L25-Gammaproteobacteria | 6732 | 26 | 139 | 19 | 36 | 18 |
| L31-Actinobacteria | 1737 | 49 | 40 | 3 | 9 | 11 |
| L31-Coriobacteria | 303 | 33 | 8 | 1 | 2 | 4 |
| L31-Corynebacteriaceae | 360 | 76 | 51 | 5 | 10 | 8 |
| L31-Firmicutes | 8296 | 32 | 18 | 0 | 6 | 6 |
| L31-Gammaproteobacteria | 8743 | 54 | 27 | 4 | 8 | 11 |

| | | | | | | |
|---|---|---|---|---|---|---|
| S4-Bacteroidia | 558 | 38 | 10 | 2 | 3 | 5 |
| S4-Clostridia | 496 | 65 | 65 | 3 | 26 | 17 |
| S4-Flavobacteria | 931 | 34 | 11 | 1 | 0 | 7 |
| S4-Fusobacteriales | 561 | 65 | 36 | 5 | 12 | 11 |
| S6:S18-Chlorobi | 67 | 78 | 31 | 4 | 14 | 6 |
| S10-Clostridia | 1294 | 86 | 41 | 4 | 19 | 13 |
| S15-Flavobacteria | 2449 | 62 | 26 | 7 | 10 | 6 |
| S15-Halobacteria | 285 | 67 | 55 | 14 | 7 | 11 |
| S15-Methanomicrobia | 194 | 79 | 49 | 21 | 10 | 7 |
| S16-Flavobacteria | 157 | 35 | 11 | 2 | 3 | 3 |

**Table S3.** Additional data on experimentally annotated transcription start sites (TSSes) and predicted r-leaders. This table presents data underlying and relating to Fig. 1. “R-leader name”: as in Fig. 1. “Organism (RefSeq sequence accession)”: the full name of the organism, and the RefSeq [31] accession of the relevant chromosome. “Citation”: a citation of the source of the TSS data. The coordinates in these papers refers to the RefSeq sequence accession given in the previous column. (In some cases, they refer to a sequence in the GenBank database [32] that is identical to the given RefSeq accession.) ”Upstream gene position”: the coordinates of the 5′ and 3′ ends of the upstream gene (which may encode a protein or a tRNA). If the 5′ coordinate (first number) is greater than the 3′ coordinate (second number), then the upstream gene is located on the reverse-complement strand of the RefSeq sequence. This information comes from the RefSeq genome annotation. “Position of TSS(es)”: list of TSS positions. This information comes from the given citation. “R-leader position”: the coordinates of the 5′ and 3′ ends of the r-leader regulating the given gene (next column). This information comes from our alignments (Additional file 2). “Regulated gene”: the coordinates of the 5′ and 3′ ends of the gene immediately downstream of the r-leader. We predicted that this gene is regulated by the r-leader. This information comes from the RefSeq genome annotation. “TSS pos.”: same meaning as in Fig. 1. These values can be calculated from the “Position of TSS(es)” and “R-leader position” columns.

| R-leader name | Organism (RefSeq sequence accession) | Citation | Upstream gene position | Position of TSS(es) | R-leader position | Regulated gene position | TSS pos. |
|---|---|---|---|---|---|---|---|
| eL15-Euryarchaeota | *Haloferax volcanii* DS2 (NC_013967.1) | [33] | 497121-496045 | 495928 | 495895-495835 | 495816-495226 | -33 |
| eL15-Euryarchaeota | *Thermococcus kodakarensis* KOD1 (NC_006624.1) | [34] | 1280610-1281680 | 1280551 | 1280545-1280502 | 1280482-1279898 | -6 |
| L20-Delta-proteobacteria | *Geobacter sulfurreducens* (NC_002939.5) | [35] | 1664628-1664825 | 1664821, 1664819, 1664816 | 1664846-1664908 | 1664915-1665268 | -25 |
| L31-Actinobacteria | *Streptomyces coelicolor* A3(2) (NC_003888.3) | [36] | 5828747-5829895 | 5829951 | 5829950-5829996 | 5830020-5830244 | +1 |
| L31-Coryne-bacteriaceae | *Coryne-bacterium glutamicum* | [37] | 928712-928398 | 928804,928836,928840 | 928877-928932 | 928944-929210 | -37 |

| | | | | | | | |
|---|---|---|---|---|---|---|---|
| | ATCC 13032 (NC_006958.1) | | | | | | |
| L31-Firmicutes | *Bacillus licheniformis* DSM 13 (NC_006322.1) | [38] | 3780008-3781291 | 3779958 | 3779923-3779890 | 3779880-3779680 | -35 |
| L31-Firmicutes | *Staphylo-coccus aureus* USA300-ISMMS1 (NC_010079.1) | [39] | 2237762-2236446 | 2236392 | 2236367-2236338 | 2236328-2236074 | -25 |
| L31-Gammaproteo-bacteria | *E. coli* (NC_000913.3) | [40] | 4126810-4124612 | 4126908, 4126911, 4126913 | 4126954-4127006 | 4127013-4127225 | -41 |
| L31-Gammaproteo-bacteria | *Shewanella oneidensis* MR-1 (NC_004347.2) | [41] | 4280501-4279464 | 4279371, 4279364 | 4279346-4279296 | 4279288-4279076 | -18 |
| S6:S18-Chlorobi | *Chlorobaculum tepidum* TLS (NC_002932.3) | [42] | 2021087-2020326 | 2020246 | 2020240-2020165 | 2020134-2019739 | -6 |
| S15-Halobacteria | *Haloferax volcanii* DS2 (NC_013967.1) | [33] | 1048559-1048630 | 1048442 | 1048444-1048382 | 1048314-1047847 | +2 |

**Table S4.** R-leaders and their operons in *E. coli* and *B. subtilis*. Based on data compiled in [12]. The column "Distinct RNA structures?" refers to whether the *E. coli* and *B. subtilis* r-leaders belong to different structural classes. Question marks indicate cases where no relevant r-leader in *B. subtilis* has been established or its ligand has not been experimentally determined. Out of eight r-leaders in *E. coli* that regulate operons containing multiple genes, five of their protein ligands are also ligands of an r-leader in *B. subtilis*, and in two of these five cases, the r-leaders are not structurally related. In the remaining three cases, no *B. subtilis* r-leader is known for any of the r-proteins encoded by the operons. The data in this table support two conclusions regarding multi-gene operons, as described in the main text. First, for r-leaders that regulate multi-gene operons, the ligand is often not encoded by the immediately downstream gene. For example, the top row shows that the immediately downstream gene of the L1-binding r-leader in *E. coli* actually encodes L11. Thus, the ligand (L1) is not encoded by the immediately downstream gene (which encodes L11). Second, this table also agrees with the hypothesis that certain r-proteins are the target of multiple r-leaders in multiple organisms, even when the r-leaders exhibit distinct structures. For example, the fifth row concerns the two r-leader motifs that bind L20, one in *E. coli* and one in *B. subtilis*. In both cases, L20 is the ligand, even though L35 is also encoded by a gene in the regulated operon.

| R-proteins encoded by operon in *E. coli* (in order of genes) | Ligand of r-leader in *E. coli* | R-proteins encoded by operon in *B. subtilis* (in order of genes) | Ligand of r-leader in *B. subtilis* | Distinct RNA structures? |
|---|---|---|---|---|
| L11, L1 | L1 | L1 | L1 | No |
| S10, L3, L4, L23, L2, S19, L22, S3, L16, L29, S17, L14, L24 | L4 | S10, L3, L4, L23, L2, S19, L22, S3, L16, L29, S17, L14, L24, L5, S14, S8, L6, L18, S5, L30, L15 | ? | ? |
| L10:L7/L12 | L10:L7/L12 | L10:L7/L12 | L10:L7/L12 | No |
| L13, S9 | L13 | | ? | ? |
| L35, L20 | L20 | L35, L20 | L20 | Yes |
| L25 | L25 | | ? | ? |
| S1 | S1 | | ? | ? |

| | | | | |
|---|---|---|---|---|
| S2 | S2 | S2 | S2 | No |
| S13, S11, S4, L17 | S4 | S4 | S4 | Yes |
| S6:S18, L9 | S6:S18 | S6:S18 | S6:S18 | No |
| S7 | S7 | | ? | ? |
| L5, S14, S8, L6, L18, S5, L30, L15 | S8 | S10, L3, L4, L23, L2, S19, L22, S3, L16, L29, S17, L14, L24, L5, S14, S8, L6, L18, S5, L30, L15 | ? | ? |
| S15 | S15 | S15 | S15 | Yes |
| S20 | S20 | | ? | ? |

**Table S5.** Association of L31 r-leader motifs with CXXC zinc-binding peptide motifs. Some r-motifs (i.e., the L31 r-leader motifs in Coriobacteria and Firmicutes) do not appear to correlate with zinc-binding. These motifs are very unlikely to play a role in zinc homeostasis. The remaining three motifs are either consistently associated with zinc-binding or consistently associated with non-zinc-binding proteins. It is possible that one or more of these three motifs is a regulator related to zinc, but we find this possibility less likely in view of the fact that the five motifs are not consistent with each other. See Supplementary Text for details.

| Motif Name | Number of regulated genes encoding complete proteins with CXXC (i.e., zinc-binding) | Number of regulated genes encoding complete proteins lacking CXXC (i.e., non-zinc-binding) | Percentage that contain CXXC |
|---|---|---|---|
| L31-Actinobacteria | 2052 | 0 | 100.0% |
| L31-Coriobacteria | 50 | 33 | 60.2% |
| L31-Corynebacteriaceae | 0 | 100 | 0.0% |
| L31-Firmicutes | 555 | 277 | 66.7% |
| L31-Gammaproteobacteria | 500 | 1 | 99.8% |

**Table S6.** Alignments from the Rfam Database [14] that were used to analyze binding sites within rRNAs.

| Domain of life | rRNA molecule | Rfam accession |
|---|---|---|
| Bacteria and Archaea | 5S | RF00001 |
| Bacteria | 16S | RF00177 |
| Bacteria | 23S | RF02541 |
| Archaea | 16S | RF01959 |
| Archaea | 23S | RF02540 |

**Table S7.** Papers used to analyze rRNA nucleotides that are known to interact with specific r-proteins. All r-proteins were analyzed using PyMol as described in Materials & Methods, but where a paper was available, we found that the PyMol results did not deviate significantly from the previously established interactions.

| R-protein | Citations |
|---|---|
| L2 | [43–45] |
| L3 | [46, 47] |
| L4 | [48, 49] |
| L6 | [50, 51] |
| L13 | None |
| L15 | [52] |
| L17 | None |
| L19 | None |
| L20 | [20, 53, 54] |
| L25 | [55, 56] |
| L31 | None |
| S4 | [57–59] |
| S6 | [24, 25] |
| S13 | [60, 61] |
| S15 | [12, 27, 62, 63] |
| S16 | [64] |

**Figure S1** (next page)**.** New r-leader motifs whose structural features are essentially the same as a previously published r-leader. Left panel: novel motifs (this study); the text "novel" refers to the alignments produced in this work. Middle panel: relevant binding sites in rRNA. Right panel: previously published r-leaders binding the same r-proteins that were experimentally validated or computationally predicted, and that have published multiple-sequence alignments. Annotations are the same as in Fig. 2 and 3. Helix numbers refer to the same source as Fig. 3. All drawings of previously predicted r-leaders in all Figures S1-S11 use alignments that were included as supplementary data of the relevant paper or are available in the Rfam Database [14] (identifying information in Table S1). (**a**) L25 r-leader in Gammaproteobacteria. Our alignment is similar to the previously published alignment [21], but has additional potential hairpins and is present in a wider variety of Gammaproteobacteria. The binding site in 5S rRNA for L25 is shown, with yellow shading as in Fig. 3. The previously published alignment was not made available in machine-readable format, and is therefore not shown. (**b**) S10 r-leader in Fusobacteria (this study). Two disjoint S10-binding regions in 16S rRNA and the L4-binding region are shown in the middle column. S10 is encoded by the immediately downstream gene, and L4 is a previously established r-leader ligand [8]. The yellow shading of the r-leader motifs shows a possible similarity with the rRNA binding site for S10, based on our analysis. (**c**) L19 r-leader in Flavobacteria. The internal loop enclosed by a C-G and a U-A base pair significantly resembles the previously published Firmicutes motif [8]. Two relevant binding sites in different parts of 16S rRNA and in 23 rRNA are shown. The figure appears on the next page.

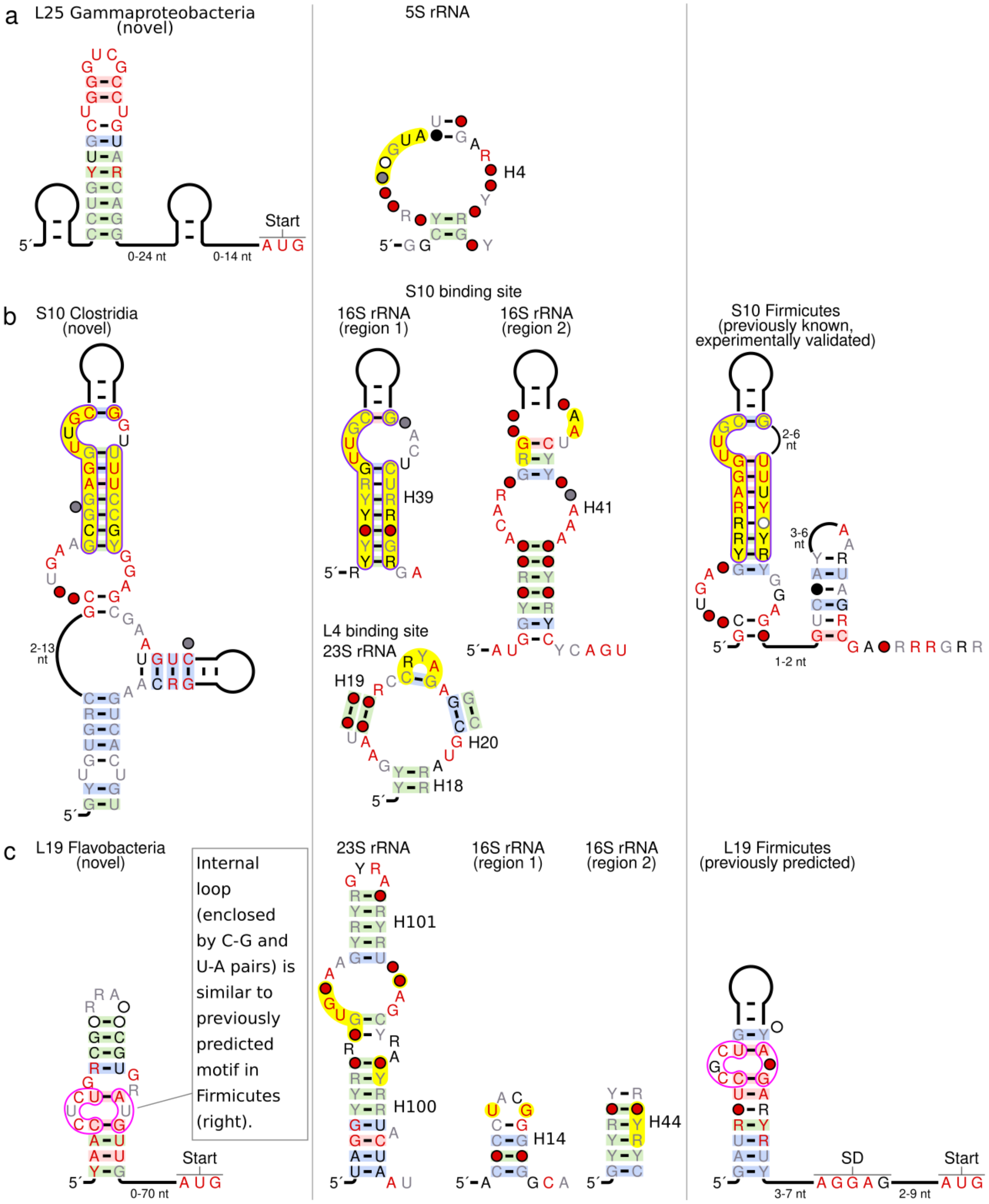
a
L25 Gammaproteobacteria (novel)
5´
0-24 nt
0-14 nt
Start
A U G
5S rRNA
H4
b
S10 Clostridia (novel)
2-13 nt
S10 binding site
16S rRNA (region 1)
H39
16S rRNA (region 2)
H41
L4 binding site 23S rRNA
H19
H20
H18
S10 Firmicutes (previously known, experimentally validated)
2-6 nt
3-6 nt
1-2 nt
c
L19 Flavobacteria (novel)
Internal loop (enclosed by C-G and U-A pairs) is similar to previously predicted motif in Firmicutes (right).
0-70 nt
Start
23S rRNA
H101
H100
16S rRNA (region 1)
H14
16S rRNA (region 2)
H44
L19 Firmicutes (previously predicted)
SD
3-7 nt
2-9 nt
Start

**Figure S2.** Novel r-leaders, rRNA and previously established r-leaders related to r-proteins L2 and L4. The left, middle and right panels have the same meaning as in Figure S1. Annotations are the same as in Figs. 2 and 3. Helix numbers refer to the same source as Figure 3.

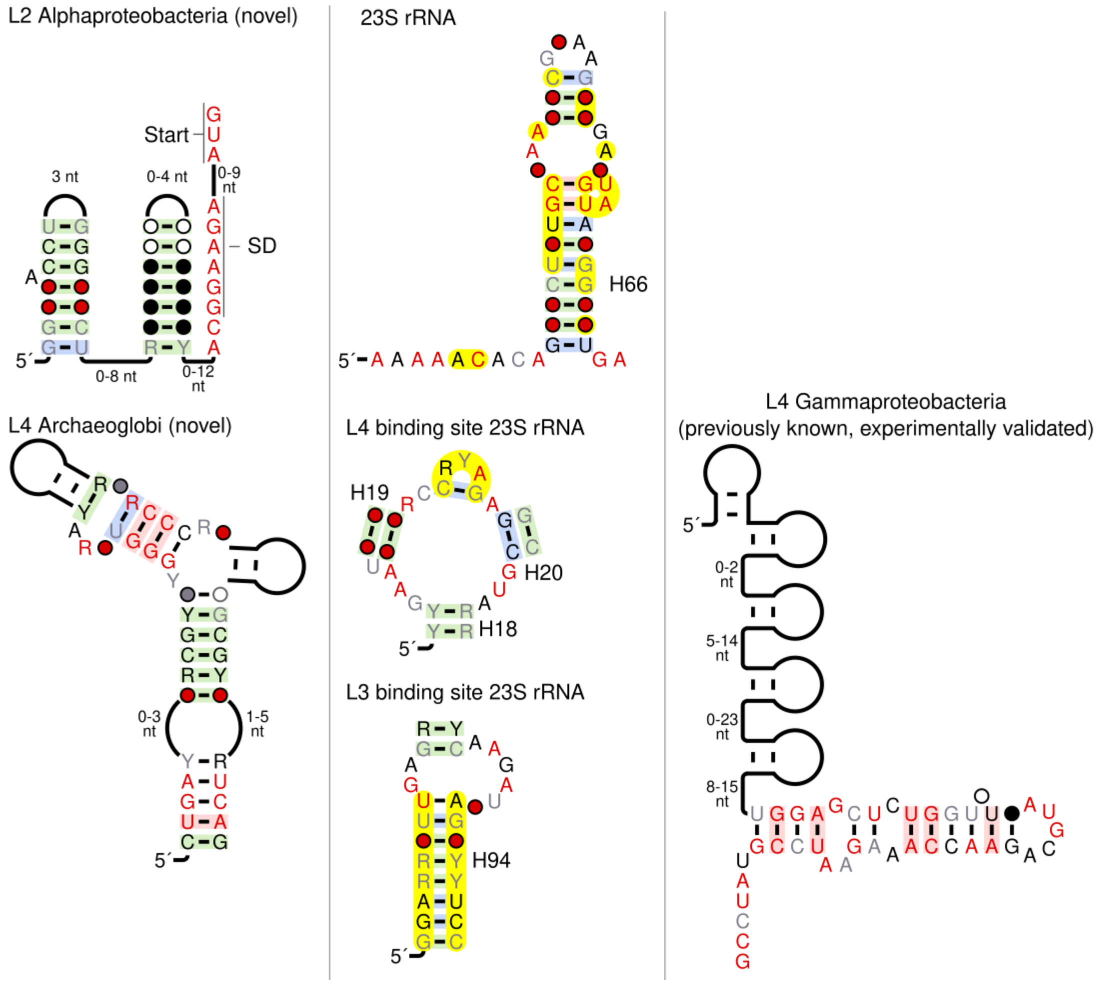

**Figure S3.** Novel r-leader, rRNA and previously established r-leaders related to r-protein L13. The left, middle and right panels have the same meaning as in Figure S1. Two disjoint regions of the rRNA make contact with the r-protein. Annotations are the same as in Figs. 2 and 3. Helix numbers refer to the same source as Fig. 3.

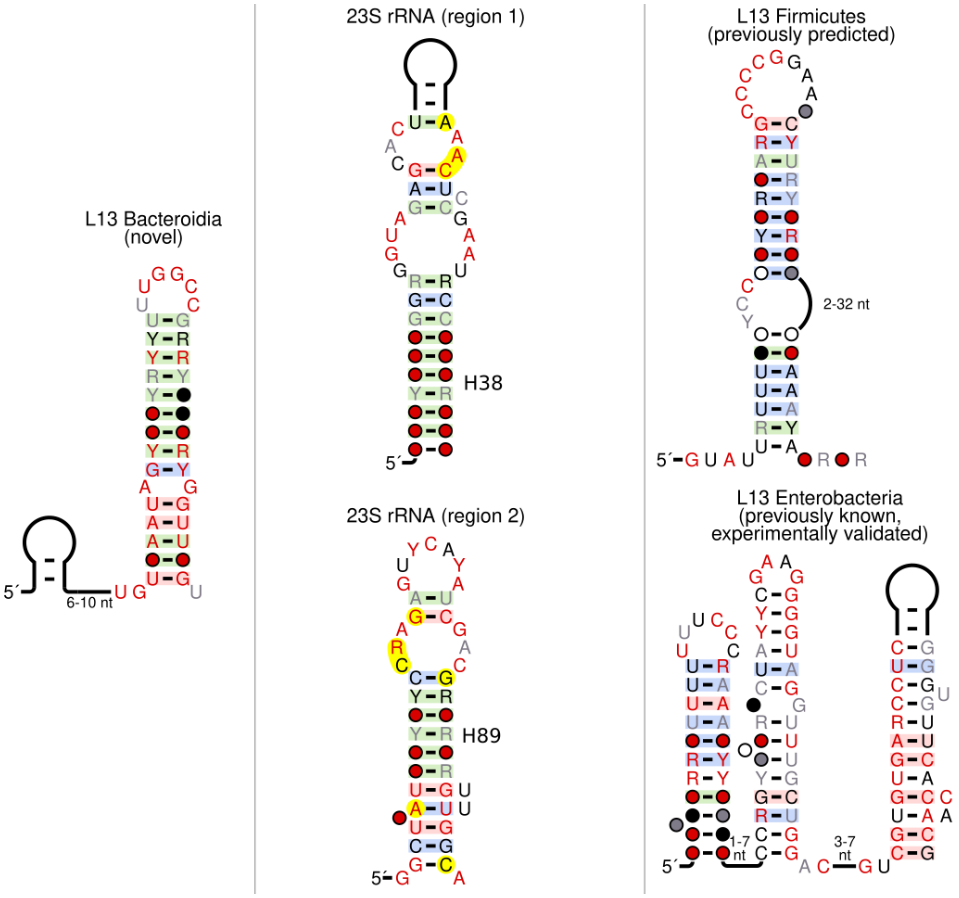

**Figure S4.** Novel r-leader, rRNA and previously established r-leaders related to r-protein eL15. The left, middle and (empty) right panels have the same meaning as in Figure S1. The right panel is empty because no eL15 r-leader has previously been established or predicted. The information in this figure is the same as in Fig. 3. We included it as a supplementary figure so that the supplementary figures are comprehensive. Annotations are the same as in Figs. 2 and 3. Helix numbers refer to the same source as Fig. 3.

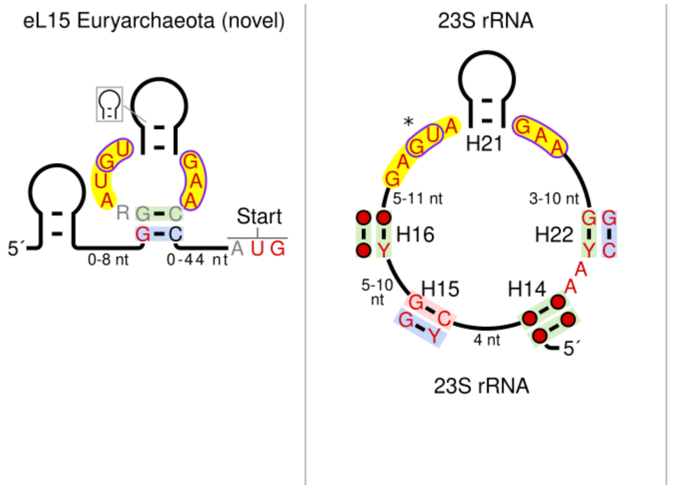

**Figure S5.** Novel r-leader, rRNA and previously established r-leaders related to r-protein L17. The left, middle and right panels have the same meaning as in Figure S1. The previously published L17 Downstream Element (L17DE) motif [19] is shown. However, if the L17DE motif does bind the L17 protein, it would function in the 3´ UTR, and therefore would not fit the strict definition of an r-leader. Annotations are the same as in Figs. 2 and 3. Helix numbers refer to the same source as Fig. 3.

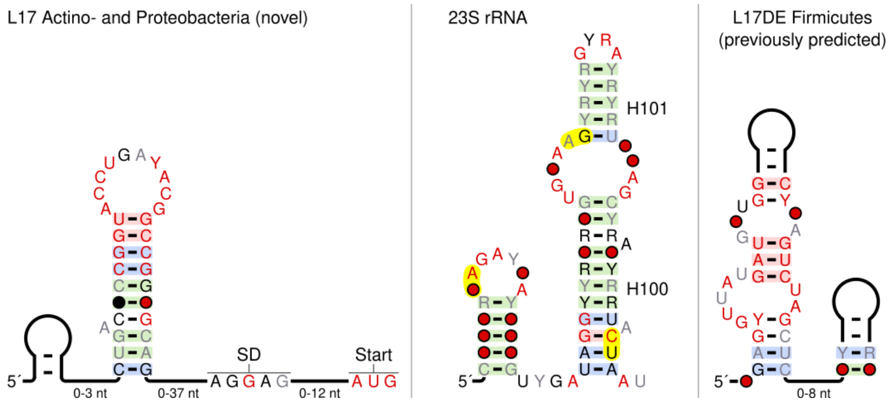

**Figure S6.** Novel r-leader, rRNA and previously established r-leaders related to r-protein L20. The left, middle and right panels have the same meaning as in Figure S1. An additional previously published L20 r-leader in Gammaproteobacteria is not shown because no alignment for it is available [12]. Annotations are the same as in Figs. 2 and 3. Helix numbers refer to the same source as Figure 3.

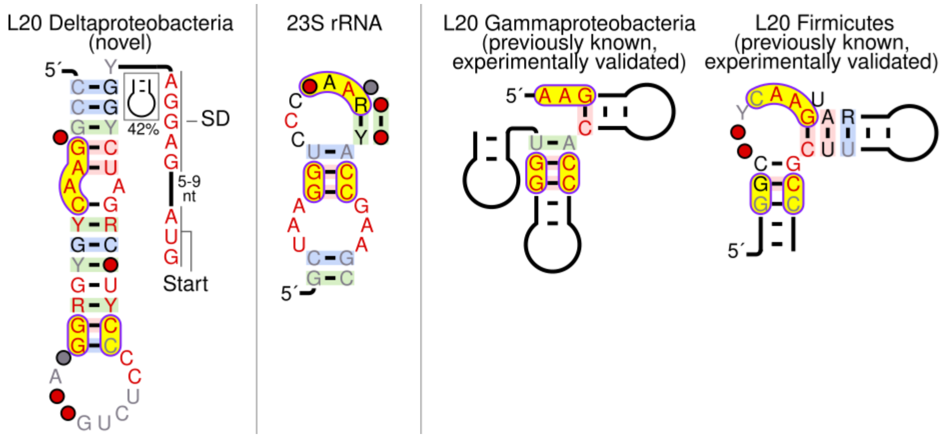

**Figure S7.** Novel r-leaders, rRNA and previously established r-leaders related to r-protein L31. The left, middle and right panels have the same meaning as in Figure S1. A version of the Firmicutes L31 motif with a conserved CUU sequence on its 5’ end is depicted. This version of the motif was not depicted in Fig. 2 because we are not persuaded that the CUU sequence is truly of biological significance (see Supplementary Text). The right panel is empty, because no r-leaders for L31 have previously been published. Annotations are the same as in Figs. 2 and 3. Helix numbers refer to the same source as Fig. 3.

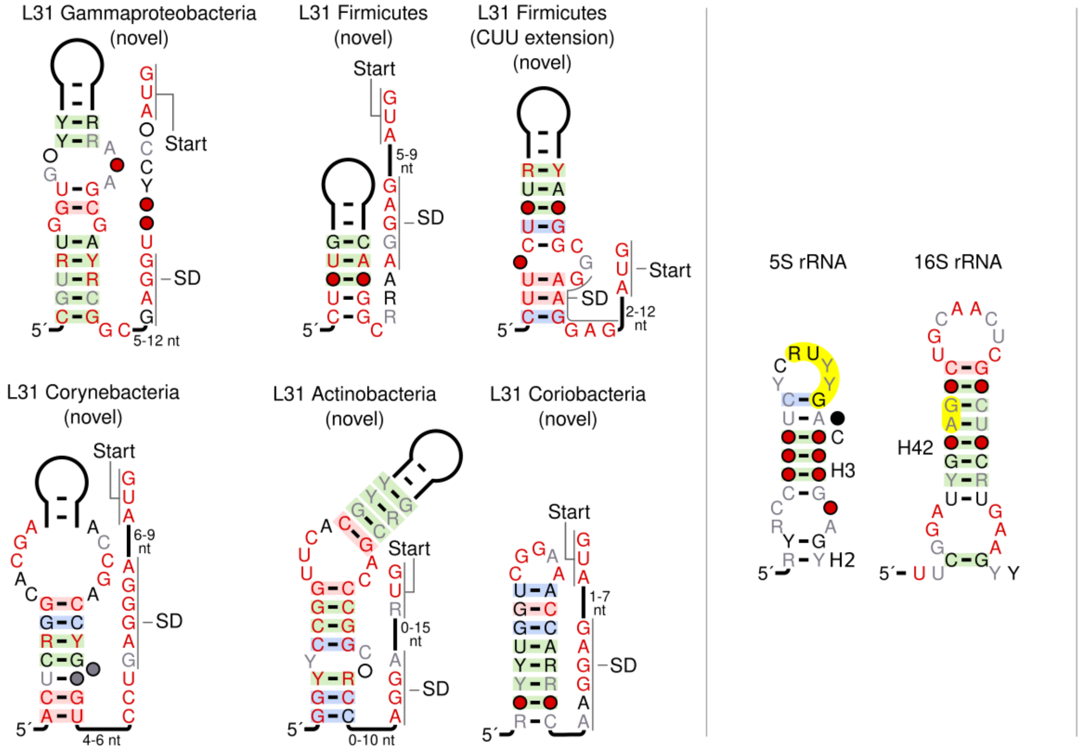

**Figure S8** (next page)**.** Novel r-leaders, rRNA and previously established r-leaders related to r-protein S4. The left, middle and right panels have the same meaning as in Figure S1. An additional region of 16S rRNA that interacts with the S4 is shown that is not depicted in Fig. 3. This region was not depicted in Fig. 3 because it does not seem similar to any of the motifs. The rRNA binding site of S13 is also shown because two of the motifs are found immediately upstream of genes that encode S13. We did not, however, find any meaningful evidence of rRNA imitation related to S13. Two versions of an alignment of S4 leaders in Firmicutes have been published. We compared our motifs to both (see text). They are labelled in the figure as "(previously known, Yao, *et al.*)" [8] and "(previously known, Deiorio-Haggar, *et al.*)" [9]. Annotations are the same as in Figs. 2 and 3. Helix numbers refer to the same source as Fig. 3. The figure appears on the next page.

S4 Fusobacteriales
(novel)

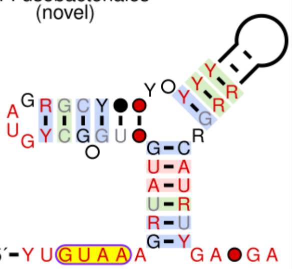

S4 Clostridia
(novel)

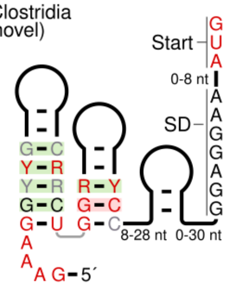


S4 Flavobacteria
(novel)

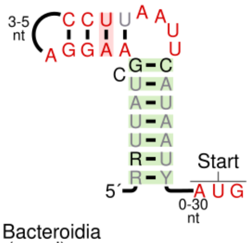


S4 Bacteroidia
(novel)

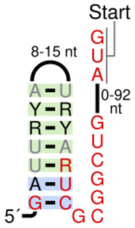


S4 binding site 16S rRNA

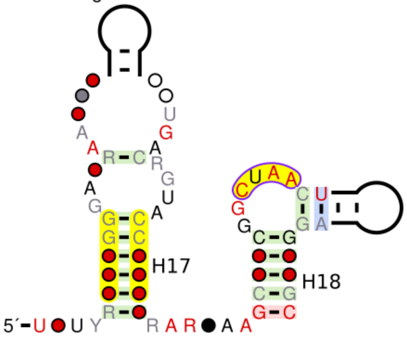


S13 binding site 16S rRNA

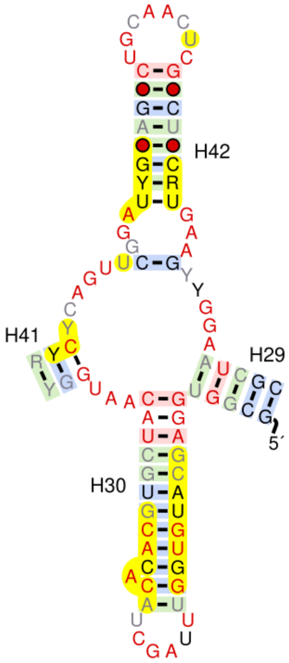


S4 Gammaproteobacteria (previously known, experimentally validated)

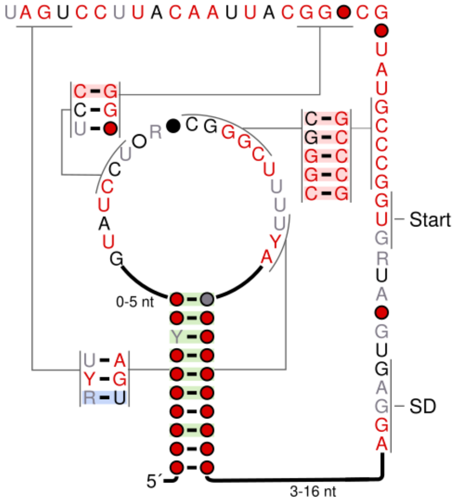


S4 Firmicutes (previously known, experimentally validated, Deiorio-Haggar *et al.)*

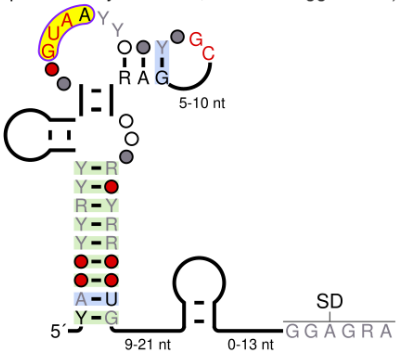


S4 Firmicutes (previously known, experimentally validated, Yao *et al.*)

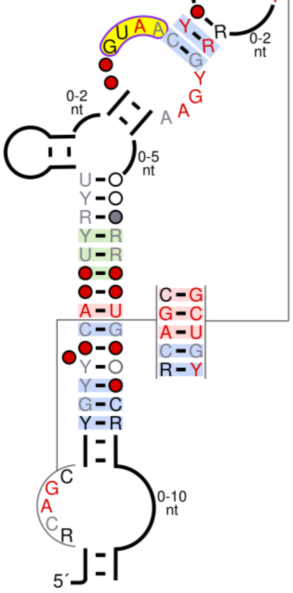

**Figure S9.** Novel r-leader, rRNA and previously established r-leaders related to r-proteins S6:S18. The left, middle and right panels have the same meaning as in Figure S1. Annotations are the same as in Figs. 2 and 3. Helix numbers refer to the same source as Figure 3.

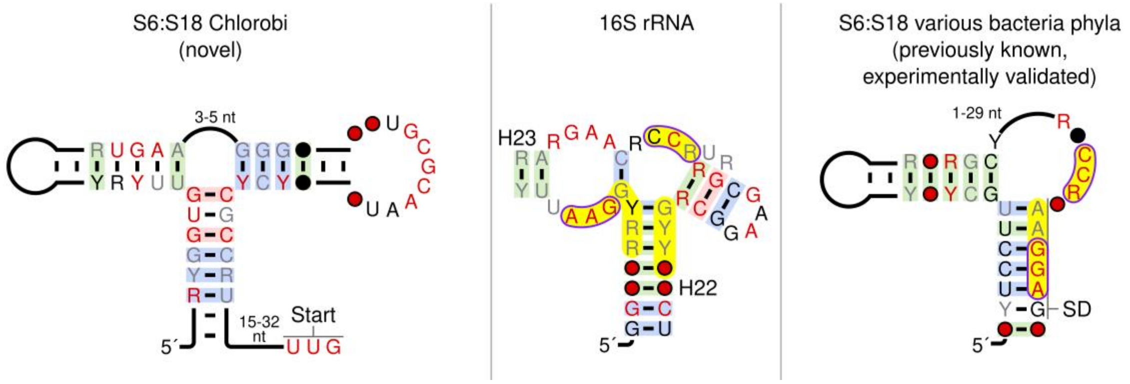

**Figure S10.** Novel r-leaders, rRNA and previously established r-leaders related to r-protein S15. The left, middle and right panels have the same meaning as in Figure S1. The putative promoter upstream of the Flavobacteria S15 r-leader motif (see Supplementary Text) is shown using DNA nucleotides (i.e., T instead of U), and there is a gap between the DNA sequence and the potential start of the RNA sequence. The hairpin depicted in cartoon form exhibits covariation, but is only present in 89% of the sequences. Archaeal and bacterial versions of the rRNA binding site are shown, due to the mixture of bacterial and archaeal motifs we found. The previously predicted S15 r-leader in *Thermus thermophilus* is not shown because no alignment is available. Annotations are the same as in Figs. 2 and 3. Helix numbers refer to the same source as Figure 3.

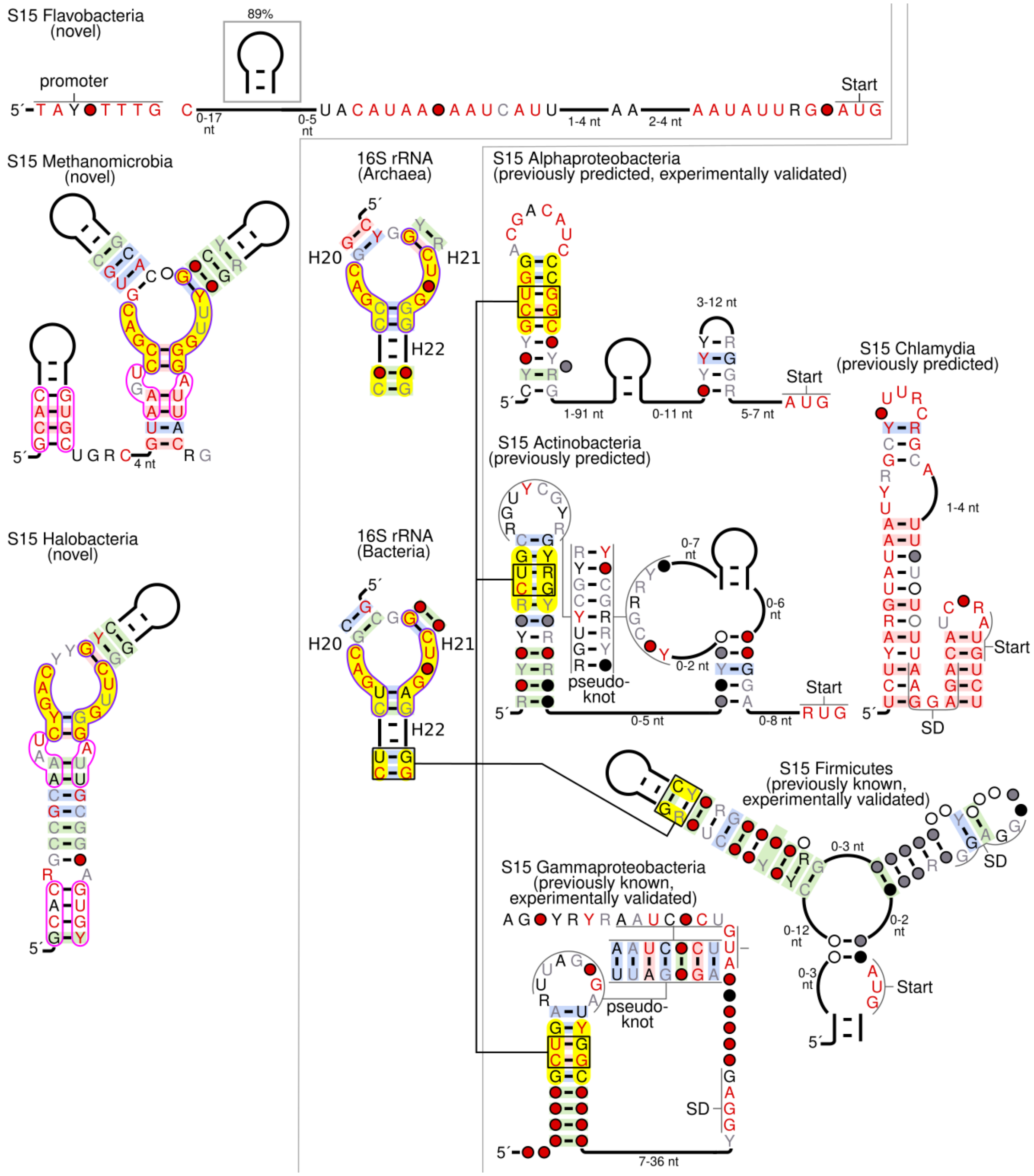
S15 Flavobacteria (novel)
89%
promoter
Start
0-17 nt
0-5 nt
1-4 nt
2-4 nt
S15 Methanomicrobia (novel)
4 nt
16S rRNA (Archaea)
H20
H21
H22
S15 Alphaproteobacteria (previously predicted, experimentally validated)
3-12 nt
1-91 nt
0-11 nt
5-7 nt
Start
S15 Chlamydia (previously predicted)
1-4 nt
Start
SD
S15 Actinobacteria (previously predicted)
0-7 nt
0-6 nt
0-2 nt
0-5 nt
0-8 nt
pseudo-knot
Start
S15 Halobacteria (novel)
16S rRNA (Bacteria)
H20
H21
H22
S15 Firmicutes (previously known, experimentally validated)
0-3 nt
0-12 nt
0-2 nt
0-3 nt
SD
Start
S15 Gammaproteobacteria (previously known, experimentally validated)
pseudo-knot
SD
7-36 nt

**Figure S11.** Novel r-leader, rRNA and previously established r-leaders related to r-protein S16. The left, middle and right panels have the same meaning as in Figure S1. No S16 r-leaders have previously been proposed. Annotations are the same as in Figs. 2 and 3. Helix numbers refer to the same source as Figure 3.

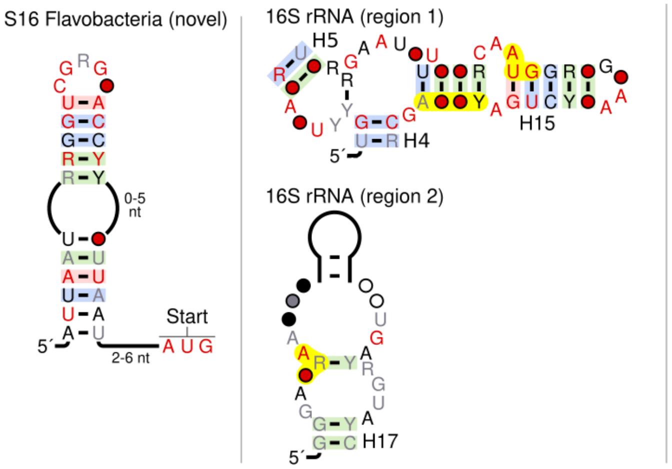
